# Investigating the “two-hit hypothesis”: effects of prenatal maternal immune activation and adolescent cannabis use on neurodevelopment in mice

**DOI:** 10.1101/2022.03.21.485178

**Authors:** Elisa Guma, Lani Cupo, Weiya Ma, Daniel Gallino, Luc Moquin, Alain Gratton, Gabriel A Devenyi, M Mallar Chakravarty

## Abstract

**Background:** Prenatal exposure to maternal immune activation (MIA) and chronic adolescent cannabis use are both risk factors for neuropsychiatric disorders. However, exposure to a single risk factor may not result in major mental illness, indicating that multiple exposures may be required for illness onset. Here, we examine whether combined exposure to prenatal MIA and adolescent delta-9-tetrahydrocannabinol (THC), the main psychoactive component of cannabis, lead to enduring neuroanatomical and behavioural changes in adulthood.

**Methods:** Mice were prenatally exposed to viral mimetic, poly I:C (5mg/kg), or vehicle at gestational day (GD)9, and postnatally exposed to chronic THC (5mg/kg, intraperitoneal) or vehicle during adolescence (postnatal day [PND]28-45). Longitudinal magnetic resonance imaging (MRI) was performed pre-treatment, PND25, post-treatment, PND50, and in adulthood, PND85, followed by behavioural tests for anxiety-like, social, and sensorimotor gating. Postmortem assessment of cannabinoid (CB)1 and 2 receptor expressing cells was performed in altered regions identified by MRI (anterior cingulate and somatosensory cortices, striatum, and hippocampus).

**Results:** Subtle deviations in neurodevelopmental trajectory and subthreshold anxiety-like behaviours were observed in mice exposed to both risk factors. Sex-dependent effects were observed in patterns of shared brain-behaviour covariation, indicative of potential sex differences in response to MIA and THC. Density of CB1 and CB2 receptor positive cells was significantly decreased in all mice exposed to MIA, THC, or both.

**Conclusions:** These findings suggest that there may be a cumulative effect of risk factor exposure on gross neuroanatomical development, and that the endocannabinoid system may be sensitive to both prenatal MIA, adolescent THC, or the combination.

## 1. Introduction

Multiple lines of evidence suggest that exposure to environmental risk factors during sensitive periods of brain development alters neurodevelopmental trajectories, increasing risk for neuropsychiatric disorders (1,2). The fetal period is thought to be especially sensitive to environmental risk factors related to neurodevelopmental disorders (3). Exposure to maternal immune activation (MIA) *in utero* has been identified as one such strong risk factor in epidemiological studies, whereby the elevation of maternal proinflammatory cytokines (and not the pathogen itself) interfere with fetal brain development (4,5). However, most maternal infections do not lead to disorders in the offspring (4). While multiple exposures may increase risk for major mental illness, MIA appears to act as a “disease primer” making individuals more susceptible to the effects of other risk factors later in life (6). In support of this hypothesis, previous work from our group found that mice prenatally exposed to MIA in early gestation (gestational day [GD]9) exhibited altered neurodevelopment, characterized by the emergence of neuroanatomical and behavioural abnormalities in the adolescent period (7), a time in which many neuropsychiatric illnesses emerge (8). However, these alterations normalized in adulthood, strongly suggesting that MIA-exposure in isolation was insufficient to create neurodevelopmental abnormalities that persisted into adulthood (7).

Cumulative exposure to drugs of abuse, particularly during the critical neurobiological remodeling that occurs during adolescence, may increase the impact of risk factors (8). Prospective, longitudinal, epidemiological evidence suggests that adolescent cannabis use may increase risk for psychosis, amongst other neuropsychiatric disorders (9–11). This is particularly evident if use begins before the age of 16 years (12). The age of sensitivity corresponds to the developmental period of the endocannabinoid system (specifically the cannabinol [CB]-1 receptor), which plays a fundamental role in brain development (13). The main psychoactive component of cannabis is delta-9-tetrahydrocannabinol (THC), which activates the endocannabinoid system via cannabinoid (CB)1 (and to some extent CB2) receptors in the brain (14). Activation of CB-1 receptors is thought to create an imbalance in excitatory-inhibitory signaling in the brain by influencing the GABA-ergic, glutamatergic, and dopaminergic systems (8). Even if cannabis exposure in adolescence increases risk for psychosis, only a minority of cannabis users develop psychosis, suggesting that cannabis use may interact with pre-existing vulnerabilities, such as prenatal MIA-exposure, and increase the risk for neuropsychiatric illness (15,16).

In this study, we leverage the strategy of performing longitudinal assays of the brain using *in vivo* magnetic resonance imaging (MRI) throughout development, and multi-behavioural testing in adulthood to investigate the potential synergistic effects of prenatal MIA-exposure in early gestation (GD9), followed by chronic adolescent THC exposure (postnatal day [PND] 28–45) in mice. Our previous work demonstrated that exposure to MIA alone leads to neuroanatomical remodeling in fetal stages (17) persisting to neurodevelopmental and behavioural abnormalities that appeared in adolescence and normalized by adulthood (7). Based on these findings, we hypothesized that exposure to each risk factor alone would result in subthreshold neuroanatomical and behavioural alterations in offspring, but that the combined exposure would lead to enduring neuroanatomical and behavioural changes in adulthood. Our neuroanatomical results partially support this hypothesis, however, investigation of CB1 and 2 receptor density suggests that there may be similar susceptibility to each risk factor.

## 2. Methods

### 2.1 Animals

Timed-mating was used to generate pregnant dams (injected intraperitoneally (i.p.) with poly I:C (POL; P1530-25MG sodium salt TLR ligand tested; Sigma Aldrich; 5mg/kg) or vehicle (SAL; 0.9% NaCl) at GD 9 (10 POL and 8 SAL dams; breeding details in **supplement 1.1**). Offspring were then randomly assigned to receive daily injections (i.p.) of THC (1:1:18 THC 5 mg/kg [Cayman Chemicals], cremophor [Sigma Aldrich], saline solution) or vehicle (1:18 cremophor:saline) throughout the adolescent period (PND28–45). The dosage was chosen to model moderate daily use, with a potency roughly equivalent to one average-sized cannabis cigarette (“joint”) with 11 % THC content, scaled to the weight of the mouse (18–20) (weight data in **supplement 1.5.1, 2.3 & supplementary figure S1**). This resulted in 4 groups: (SAL-SAL, SAL-THC, POL-SAL, POL-THC). In a separate group of dams, the immunostimulatory potential of our poly I:C was confirmed (n=4 GD9-POL, n=5 GD9-SAL; **supplement 1.2 for methods, 2.1 and supplementary table 1 for results**).

We validated our physiologically compatible THC solution (1:1:18 THC:cremophor:saline at 2.5, 5, or 10 mg/kg of THC; Cayman Chemicals, Ann Arbor, MI, USA) by measuring plasma levels of THC and two THC metabolites (11-hydroxy-delta-9-THC and 11nor-9carboxy delta-9-THC) in a separate cohort of mice using gas-chromatography mass spectrometry (GCMS) (**supplement 1.3 for methods, 2.2 and supplementary table 2 for results**). A license to possess cannabis for research purposes (LIC-ZULO6PJ7NH-2019) and an import permit (IMP-0257-2019) to purchase cannabis were obtained from Health Canada. All procedures were approved by McGill University’s Animal Care Committee under the guidelines of the Canadian Council on Animal Care.

### 2.4 Magnetic resonance imaging

#### 2.4.1 Acquisition and image processing

Longitudinal T1-weighted structural MRIs were acquired *in vivo* at postnatal day (PND) 24-26 (~childhood; pretreatment), 47-51 (~adolescence; post-treatment), 80-86 (~adulthood) (21,22) in anesthetized offspring (7,23–26). Anesthesia was induced with 3% isoflurane in oxygen and a (0.075 mg/kg bolus) dexmedetomidine injection. Anesthesia was maintained during the scan between 1.5-0.5% isoflurane, and a constant infusion of dexmedetomidine (0.05mg/kg/h continuous mg/kg during scan). Scans were conducted in a 7 Tesla Bruker, 30 cm bore magnet with AVANCE electronics, using a cryogenically cooled surface coil. A 3D FLASH (Fast, Low Angle SHot) sequence was used with a flip angle of 20 degrees and TE/TR of 4.5/20 ms (2 averages, ~14 minutes). Voxel size was 100 μm, isotropic, in a matrix of 180 by 160 by 90. Structural images were visually inspected for quality (3 removed due to motion or hydrocephalus, yielding n=242 images). Images were processed using a longitudinal two-level deformation based morphometry technique using the ANTs toolkit (http://stnava.github.io/ANTs/) for linear and non-linear registration (27) as described in our previous work (28) (**Supplement 1.4**).

**Table 1.** Sample size per timepoint following quality control. Postnatal day (PND); poly I:C (POL); saline (SAL); delta-9-tetrahidrocannabinol (THC); male (M); female (F).

|  | PND 25 |  | PND 50 |  | PND 85 |  |  |
| --- | --- | --- | --- | --- | --- | --- | --- |
|  | Males | Females | Males | Females | Males | Females | Litters |
| **SAL-SAL** | 10 | 9 | 10 | 9 | 10 | 9 | 7 |
| **POL-SAL** | 10 | 10 | 11 | 10 | 11 | 10 | 9 |
| **SAL-THC** | 10 | 8 | 10 | 8 | 9 | 9 | 7 |
| **POL-THC** | 12 | 9 | 11 | 11 | 11 | 11 | 8 |

#### 2.4.2 Statistical Analyses

A voxel-wise linear mixed-effects model (R-3.5.1, RMINC-1.5.2.2, lme4 1.1-21) was used to examine a group by age (age as a natural second order spline) interaction covarying for sex (with subject and litter as random intercepts). The False Discovery Rate (FDR)(29) correction was applied to control for multiple comparisons across voxels. First, we examined whether each risk factor alone had significant effects on neurodevelopment. We performed pairwise comparisons to determine the impact of individual or combined risk factors while holding SAL-SAL as the reference group. Given evidence of sex-differences in response to MIA and THC, a sex-by group-by-age interaction was investigated post-hoc (**Supplement 1.6.2**) (30,31).

### 2.5 Behavioural testing

#### 2.5.1 Data acquisition

Following the final scan at PND ~85, testing was performed to assay several behaviours relevant to neurodevelopmental and neuropsychiatric disorders. These included the open field test to assess exploratory and anxiety-like behaviours, wherein the distance traveled in the anxiogenic center zone was examined relative to the total distance traveled. Next, the three chambered social task was performed to assay both social preference (i.e., preference for a novel mouse relative to a nonsocial object), and social novelty (i.e., preference for a novel mouse relative to a familiar mouse) behaviours. Finally, sensorimotor gating to an acoustic startle was assessed with the prepulse inhibition (PPI) task, wherein a prepulse tone is presented (at various levels, across multiple trials) to determine whether it is successful at dampening the startle reaction to the acoustic stimulation. A 2-day resting phase was allowed between tests; details for all behavioural tests are provided in **supplement 1.5**.

#### 2.5.2 Statistical Analyses

We used linear mixed-effects models for adult behavioural data, with group and sex as fixed effects, and litter as a random intercept. A Bonferroni correction was applied (4 tests: α= 0.05/4 = 0.0125 set as significance threshold, uncorrected p-values, and corrected p-values reported). Sex differences were investigated post-hoc (**supplement 1.6.2**).

### 2.6 Partial Least Squares Analysis

To assess brain-behaviour relationships in our mice, we used a partial least squares (PLS) analysis. This multivariate technique allows us to relate two sets of variables by finding the optimal weighted linear combinations of variables that maximally covary with each other (32–34) (details in **supplement 1.7**). We performed two analyses: for the first analysis we used the final MRI timepoint cross-sectional volume data (PND 85; brain matrix); tor the second analysis we calculated the within-subject voxel-wise brain volume change from PND 50 to 85 (i.e. immediately post-treatment, to the final timepoint; brain matrix; https://github.com/CoBrALab/documentation/wiki/Create-a-nifti-of-within-subject-change-(2-timepoints,-output-from-dbm) to use as the input brain data. Both were evaluated against behavioural metrics derived from the three tests described above (**2.5**).

### 2.7 Immunohistochemistry

Two days following the last behavioural test, mice were perfused (as described in **Supplement 1.8.1**), brains were extracted, sucrose protected, and sectioned coronally at 50 μm-thickness in preparation for immunofluorescent staining [SAL-SAL (n=5), POL-SAL (n=6), POL-THC (n=6) and SAL-THC (n=6)]. Regions of interest, selected based on MRI results and MIA/THC literature, included the primary somatosensory cortex and striatum, two regions that do not show normalization in adulthood in the POL-THC group, and have been associated with sensitivity to these risk factors (7,35). Additionally, we investigated the anterior cingulate cortex (ACC), which shows post treatment effects and a normalization in adulthood. Finally, we focused on the hippocampus, a region shown to be affected following MIA-exposure by our group and others, as well as by THC exposure (7,36–38).

We examined CB1- and CB2-immunoreactive (IR) cells in our ROIs and captured average optical intensity of axon terminals per counting area (Fiji ImageJ (39); values ranged from 1-256) in all regions of interest (striatum Bregma +0.38 mm, ACC Bregma −0.94 mm, somatosensory cortex Bregma −0.94 mm, dentate gyrus (DG) and CA1 molecular layers of the hippocampus Bregma −2.80 mm). In all regions (except the striatum and CA1), cell (neuron) density (cell number/area) as well as fiber density (optical intensity/area) were quantified (details in **Supplement 1.8.2**). Although we did not costain for a neuron marker, such as NeuN, based on the morphology of stained cells and the location it is likely that the cells stained were neurons (ex: pyramidal shape in dentate gyrus) (40,41). Differences in CB1 - and CB2-IR cell (neuron) density and/or fiber density due to pre- and postnatal exposures were assessed with a two-way ANCOVA, with a prenatal [SAL/POL] by adolescent [SAL/THC] treatment interaction (sex [M/F] and hemisphere [L/R] were included as covariates). Post-hoc, pairwise testing was performed using Tukey HSD (significance level was set at *p* < 0.05).

## 3. Results

### 3.1. Alterations in neurodevelopmental trajectories

#### 3.1.1 Prenatal MIA-exposure

Overall differences between groups are summarized in **supplementary figure S2 & supplement 2.4**.Effects of MIA exposure on offspring development were assessed by comparing POL-SAL to SAL-SAL. The group by age interaction modeled with a second-order natural spline (ns) (*group:ns*(*age,2*)*1*) interaction was significant in a limited number of voxels in the cerebellum, at a lenient 20% FDR threshold (t=4.662; **Figure 2AB**). To determine whether some of the changes we observed in our previous work (7) were recapitulated here at a subthreshold level, we investigated changes in trajectory at a threshold of uncorrected p<0.01. At this exploratory level, we observed that POL exposed offspring had a smaller volume at PND 25, relative to saline, which overshot (was relatively larger) at PND 50, and then normalized at PND 85 (7). This was observed in regions such as the lateral septum, striatum, hippocampus, and somatosensory cortex. Post-hoc investigation of sex differences revealed no effects (<20%FDR).

**Figure 1.**
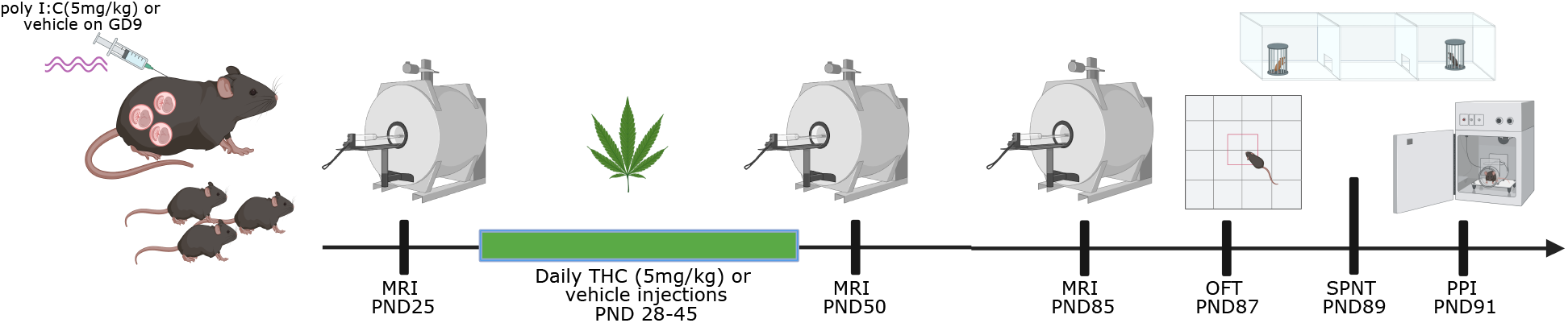
Experimental timeline. Pregnant dams were injected (i.p.) with poly I:C (5mg/kg) or vehicle (0.9% sterile NaCl solution) at gestational day (GD) 9. Offspring were weaned and sexed on postnatal day (PND) 21. Longitudinal structural magnetic resonance imaging (MRI) was performed at PND 25, 50, and 85 (denoted by the MRI image). Following the first scan on PND 25, offspring were randomly assigned to receive daily injections (i.p.) of delta-9-tetrahydrocannabinol (THC; 5 mg/kg) or vehicle (1:18 cremophor:saline) from PND 28-45. Two days following the final scan on PND85, mice were assessed in the open field test (OFT), the social preference/novelty test (SNPT), and the prepulse inhibition task (PPI), with a 2-day rest between tests.

**Figure 2.**
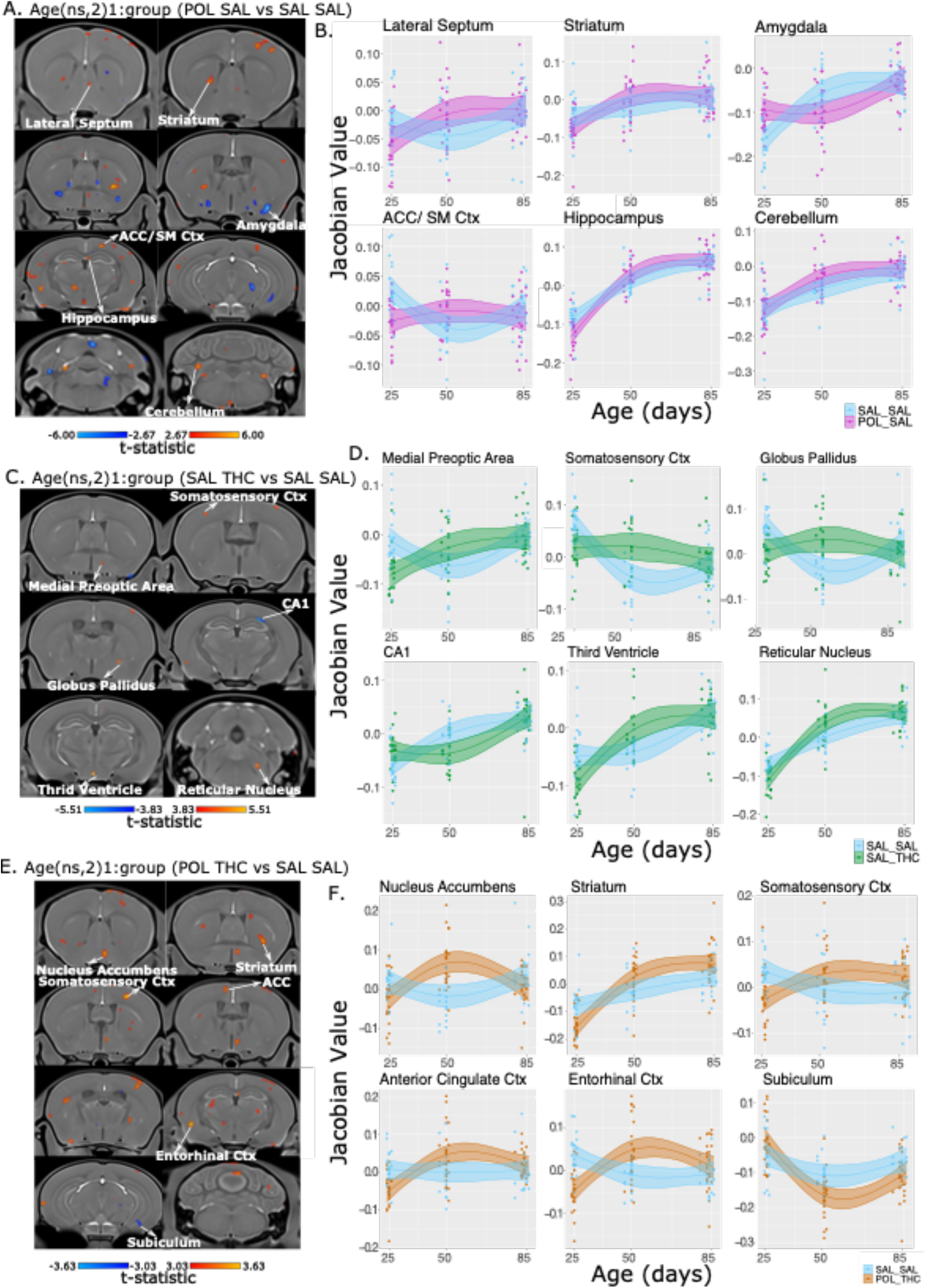
Neuroanatomical alterations due to pre- and/or postnatal risk factor exposure from preadolescence to adulthood. **A**. Exploration of neuroanatomical alteration due to prenatal MIA-exposure alone at an uncorrected threshold (p<0.01). T-statistic map of group (POL-SAL vs SAL-SAL) by age (first order natural spline (ns) of age) (t=2.67, p=0.01). **B**. Plot of peak voxels selected from regions of interest highlighted in **A**, wherein age is plotted on the x-axis, and the absolute Jacobian determinants plotted on the y-axis. Here a value of 1 means the voxel is no different than the average, anything above one is relatively larger, and below 0 is relatively smaller. Trajectories reflect the statistical model used (see **2.4.2**). **C**.Neuroanatomical alteration due to adolescent THC-exposure. T-statistic map of group (SAL-THC vs SAL-SAL) by age (first order natural spline of age) thresholded between 5% FDR (top, t=5.51) and 20% FDR (bottom, t=3.83). **D**. Plot of peak voxels selected from regions of interest highlighted in **C**, wherein age is plotted on the x-axis, and the absolute Jacobian determinants plotted on the y-axis. Trajectories reflect the statistical model used (see **2.4.2**). **E**. Neuroanatomical alteration due to combined prenatal MIA-exposure and adolescent THC-exposure. t-statistic map of group (SAL-THC vs SAL-SAL) by age (first order natural spline of age) thresholded between 5% FDR (top, t=3.63) and 10% FDR (bottom, t=3.03). **F**. Plot of peak voxels selected from regions of interest highlighted in **E**, wherein age is plotted on the x-axis, and the absolute Jacobian determinants plotted on the y-axis. Trajectories reflect the statistical model used (see **2.4.2**).

#### 3.1.2 THC

We observed a significant group by age (*group:ns*(*age,2)1*) interaction (t=5.16, 5%FDR; **Figure 2CD**) between SAL-SAL and SAL-THC groups. To evaluate more subtle neuroanatomical changes, brain maps were investigated at a more lenient threshold of 20% FDR. We observed the greatest deviations in SAL-THC mouse trajectory relative to

SAL-SAL at the PND 50, or post-treatment timepoint. This suggests that the chronic THC exposure was inducing some brain remodeling, however, we see that the trajectories normalize in later adulthood (PND 85), which may indicate that the brain had a chance to self-correct following a washout period from THC. In the medial preoptic area, somatosensory cortex, globus pallidus, third ventricle, and reticular nucleus, the SAL-THC offspring had larger volumes at PND50, whereas in the CA1 region of the hippocampus, they had smaller volumes. Post-hoc investigation of sex differences revealed no significant sex by group by age interactions.

#### 3.1.3 Combined MIA-THC

A significant group by age (*group:ns(age,2)1*) interaction (t=3.63, 5%FDR) was observed when comparing POL-THC to SAL-SAL trajectories. Again, we observed a significant deviation in trajectory in the post-treatment, PND50 timepoint wherein increased volume in the POL-THC group was observed in the nucleus accumbens, striatum, somatosensory, anterior cingulate, and entorhinal cortices, while decreases were observed in the subiculum. Interestingly, this deviation in trajectory normalized in several regions, including the nucleus accumbens, anterior cingulate and entorhinal cortices, while the deviation was sustained in later adulthood (PND85) in the striatum, somatosensory cortex, and subiculum. This suggests that some regions may recover following combined exposure, while others remain affected throughout development (**Figure 2 EF**). Investigation of the effects of POL-THC relative to POL-SAL are summarized in **supplement 2.5 and supplementary figure S3**. Post-hoc investigation of sex differences in this group revealed a significant three way interaction (*group:ns*(*age,2*)*1:sex*) (t=4.238, 5%FDR) (described in **supplement 2.7.1, supplementary figure S5**).

### 3.4 Alteration in behavior

Overall, none of the behavioural effects observed survived multiple comparisons correction. No significant effects due to prenatal MIA exposure (POL-SAL vs. SAL-SAL) or THC exposure (SAL-THC vs. SAL-SAL) were observed in the open field test, PPI, or social preference or novelty behaviours (q>0.0125). Finally, for the combined exposure group (POL-THC vs. SAL-SAL), a subthreshold increase in the relative distance traveled in the center zone of the open field was observed (t=2.344, p=0.041, q=0.164). No significant effects were observed in sensorimotor gating overall, however, when investigating effects over increasing prepulse level, there was a subthreshold interaction (t=2.239, p=0.027, q=0.108) wherein POL-THC offspring were impaired at across prepulse tones relative to SAL-SAL offspring. Again, no social deficits were observed for either preference or novelty (**Supplementary figure 4**).Finally, investigating THC effects within the prenatal POL groups (POL-SAL vs. POL-THC) revealed no significant differences, other than similar relationships in PPI reported above (t=2.239, p=0.027, q=0.108). Comparison between POL-THC and POL-SAL detailed in **Supplement 2.6.1, Supplementary figure S4, Supplementary table 3**). Behavioural results are summarized in **Supplementary table 3**. No significant sex-by-group interactions were observed (**Supplement 2.6.2, Supplementary figure S6; supplementary table 4**).

### 3.5 Brain-behaviour covariation

#### 3.5.1 PND85 brain-behaviour

Using PLS to examine adult, whole-brain, voxel-wise volume differences and 18 behavioural metrics across our 3 main tests (including sex and litter size), we identified one significant LV (p<0.00001, percent-covariance=35%; **Figure 3A**). The observed pattern identified increased pancortical, striatal, thalamic, and cerebellar volumes to be associated with decreased interactions in the social preference test, decreased sensorimotor gating abilities, highly expressed in females **(Figure 3BC**). Correlations between brain-behaviour scores suggests that all female offspring heavily express the brain-behaviour pattern, as do the SAL-SAL male offspring, but all male offspring exposed to either a single (prenatal MIA, adolescent THC) or combined risk factors (both MIA and THC) do not express the pattern. This is indicative of potential sex differences in response to risk factor exposure, with greater group differences in male offspring (**Figure 3D**).

**Figure 3.**
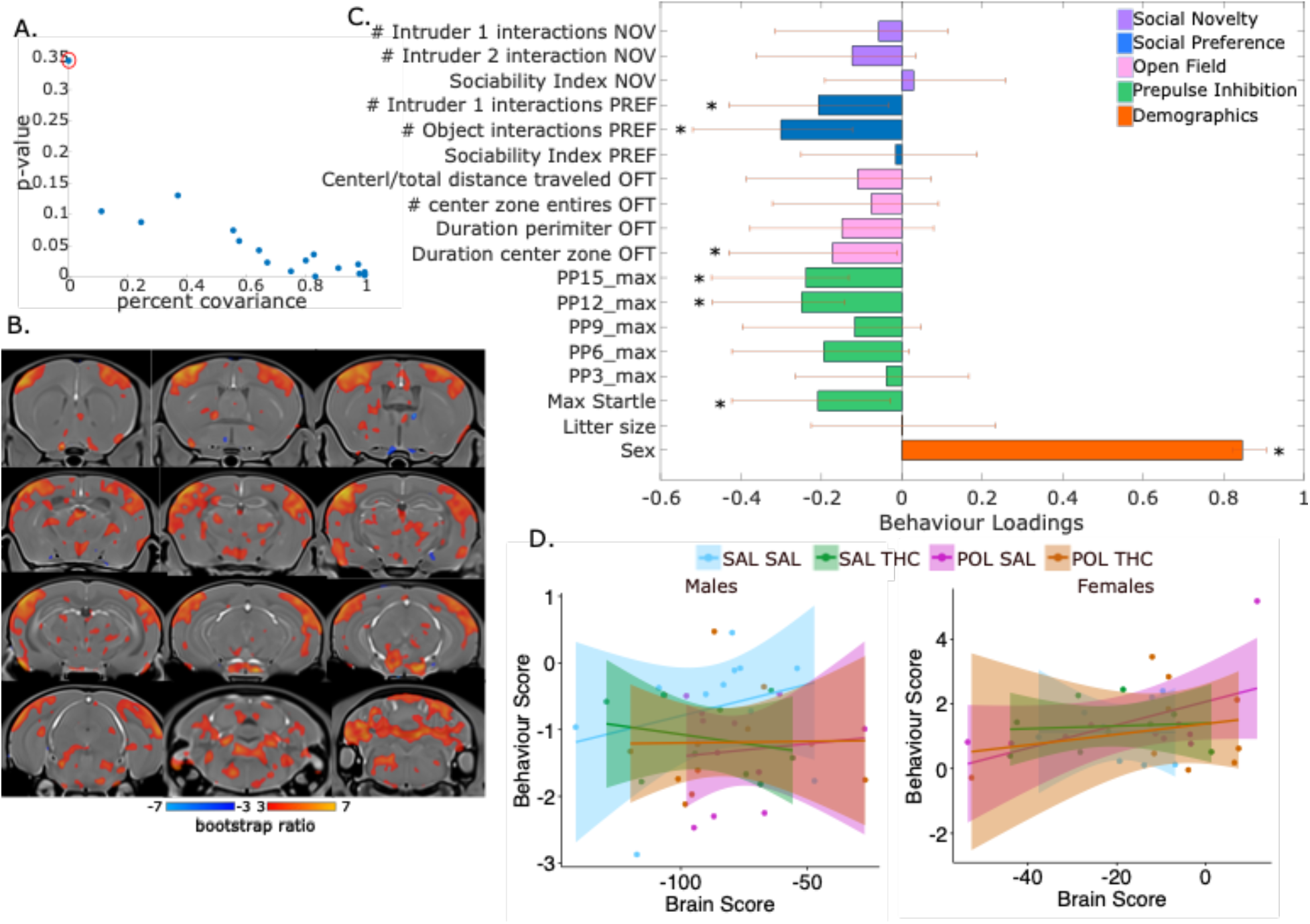
Partial least squares (PLS) analysis results for the first latent variable (LV1) comparing adult (PND85) brain and behaviour. **A**. Covariance explained (y-axis) and permutation p-values (x-axis) for all 18 LVs in the PLS analysis. LV1 is circled in red (p<0.00001, % covariance=35%) and was chosen for subsequent investigation based on the covariance explained and behavioural relevance of results. **B**. Brain loading bootstrap ratios for the LV1 deformation pattern overlaid on the population average, with positive bootstrap ratios in orange-yellow (indicative of larger volume), and negative in blue (indicative of smaller volume). Colored voxels make significant contributions to LV1. **C**. Behaviour weight for each behavioural measure included in the analysis showing how much they contribute to the pattern of LV1. Singular value decomposition estimates the size of the bars whereas confidence intervals are estimated by bootstrapping. Bars with error bars that cross the 0 line should not be considered significant, marked by *. **D**. Correlation of individual mouse brain and behaviour scores, colour coded by treatment group with a trend line per group split by sex. Male SAL-SAL offspring, and all female offspring express the pattern more strongly; male offspring exposed to any of the risk factors express the pattern less strongly. SAL-SAL (cyan), POL-SAL (magenta), SAL-THC (green), POL-THC (orange).

#### 3.5.2 Association between volume change from post-treatment to adulthood and adult behaviour

Next, we used PLS to examine within-subject volume change (value representing difference in volume between timepoints) from the post-treatment timepoint (PND50) to the adult timepoint (PND85) at a voxel level across the brain and the same 18 behavioural and demographic metrics as in **3.5.1**. We identified two significant LVs (LV1: p<0.00001, %covariance=24%; LV2: p=0.01, %covariance=19%; **Supplementary figure S7 & supplement 2.7**). For LV1, the observed pattern identified increased somatomotor, striatal, hippocampal, and cerebellar volumes to be associated with fewer interactions in the social preference test, greater impairments in sensorimotor gating, and to be more expressed in females, similar to the pattern observed in **3.5.1 (Figure 4AB**). Correlations between brain-behaviour scores show that females express this pattern, while males show some group differences, with greater loading in SAL-SAL and POL-THC groups (left two plots **Figure 4C**). Removal of outliers (right two plots **Figure 4C**) suggests that male offspring express this pattern, as do females, except for the SAL-THC female offspring, who seem to express this pattern less, indicative of potential sex differences.

**Figure 4.**
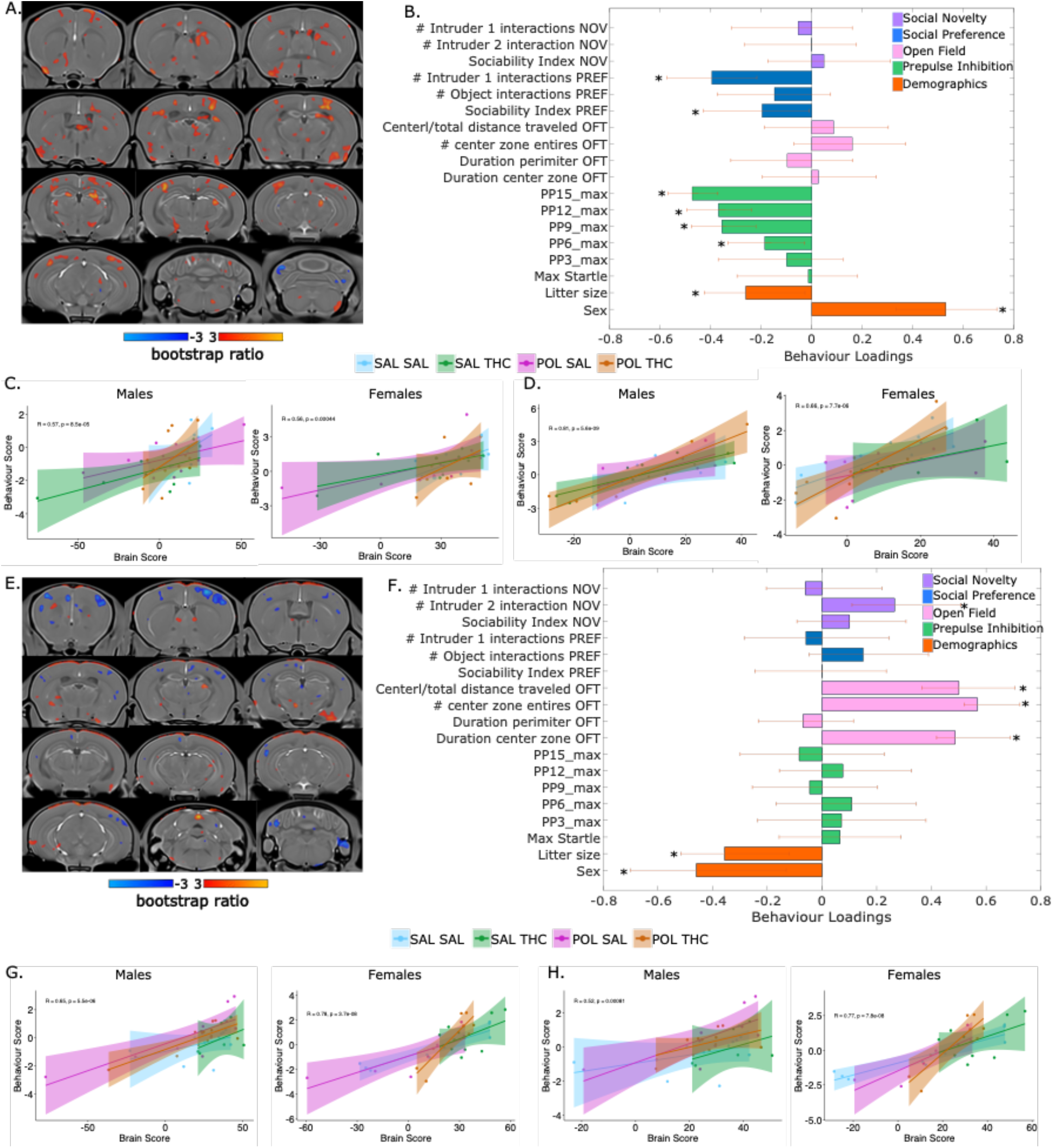
Partial least squares (PLS) analysis results for the first and second latent variables (LV) comparing the difference between PND50 and PND85 voxel-wise brain volume, and behaviour. **A**. Brain loading bootstrap ratios for the LV1 deformation pattern overlaid on the population average, with positive bootstrap ratios in orange-yellow (indicative of positive change in volume), and negative in blue (indicative of negative change in volume). Colored voxels make significant contributions to LV1. **B**. Behaviour weight for each behavioural measure included in the analysis showing how much they contribute to the pattern of LV1. Singular value decomposition estimates the size of the bars whereas confidence intervals are estimated by bootstrapping. Bars with error bars that cross the 0 line should not be considered significant and are marked by *. **C**. Correlation of individual mouse brain and behaviour scores, color coded by treatment group with a trend line per group split by sex. **D**. Correlation of individual brain-behaviour scores with the exclusion of outliers (determined using the following cut offs: minimum (quartile 1 - 1.5x interquartile range) and maximum (quartile3 + 1.5x interquartile range). **E**. Brain loadings for LV2 (as in **A**). **F**. Behaviour loadings for LV2. **G**. Correlation for individual mouse brain and behaviour score, coloured by treatment and split by sex. **H**. Correlation of individual brain-behaviour scores with the exclusion of outliers. SAL-SAL (cyan), POL-SAL (magenta), SAL-THC (green), POL-THC (orange).

The pattern captured by LV2 reflects decreased cortical (somatomotor) and cerebellar volume and increased thalamic volume to associate with increased social novelty interactions, increased locomotion in the open field test, and to be more associated with male mice from smaller litters (**Figure 4EF**). Brain-behaviour score correlations suggest that all offspring express this pattern, with a potential difference in the POL-THC exposed female offspring (left two plots **Figure 4G**). Removal of outliers (right two plots **Figure 4G**) confirms this association.

### 3.6 CB1- and CB2-IR neurons and axonal fiber and density are decreased across all brain regions due to exposure to any risk factor

Neuron cell density and axonal fiber density for CB1- and CB2-IR cells were significantly reduced across all brain regions for any of the three risk-factor exposed groups, relative to the SAL-SAL controls. A significant pre-by postnatal treatment interaction was observed for CB1-IR (F(5,40)=12.651, p=0.00098) and CB2-IR density (F(5,40)=6.051, p=0.01832). Similar effects were observed in the somatosensory cortex for CB1-(F(5,40)=41.537, p=1e-06) and CB2-IR density (F(5,40)=14.823, p=0.000416). In both these regions, Tukey’s post-hoc pairwise testing confirmed that POL-SAL, SAL-THC, and POL-THC all had significantly decreased CB1- and CB2-IR density relative to the SAL-SAL controls (p>0.05). In the striatum, the same pattern was observed for fiber density measures (cell density not quantified), with a significant pre-by postnatal treatment interaction for both CB1(F(5,40)=4.632, p=3.80e-05) and CB2-IR (F(5,40)=4.478)). As with the ACC and somatosensory cortex, in the striatum, all three risk factor groups also had significantly reduced CB1-IR and CB2-IR fiber density relative to the SAL-SAL controls (p<0.01) (**Figure 5**).

**Figure 5.**
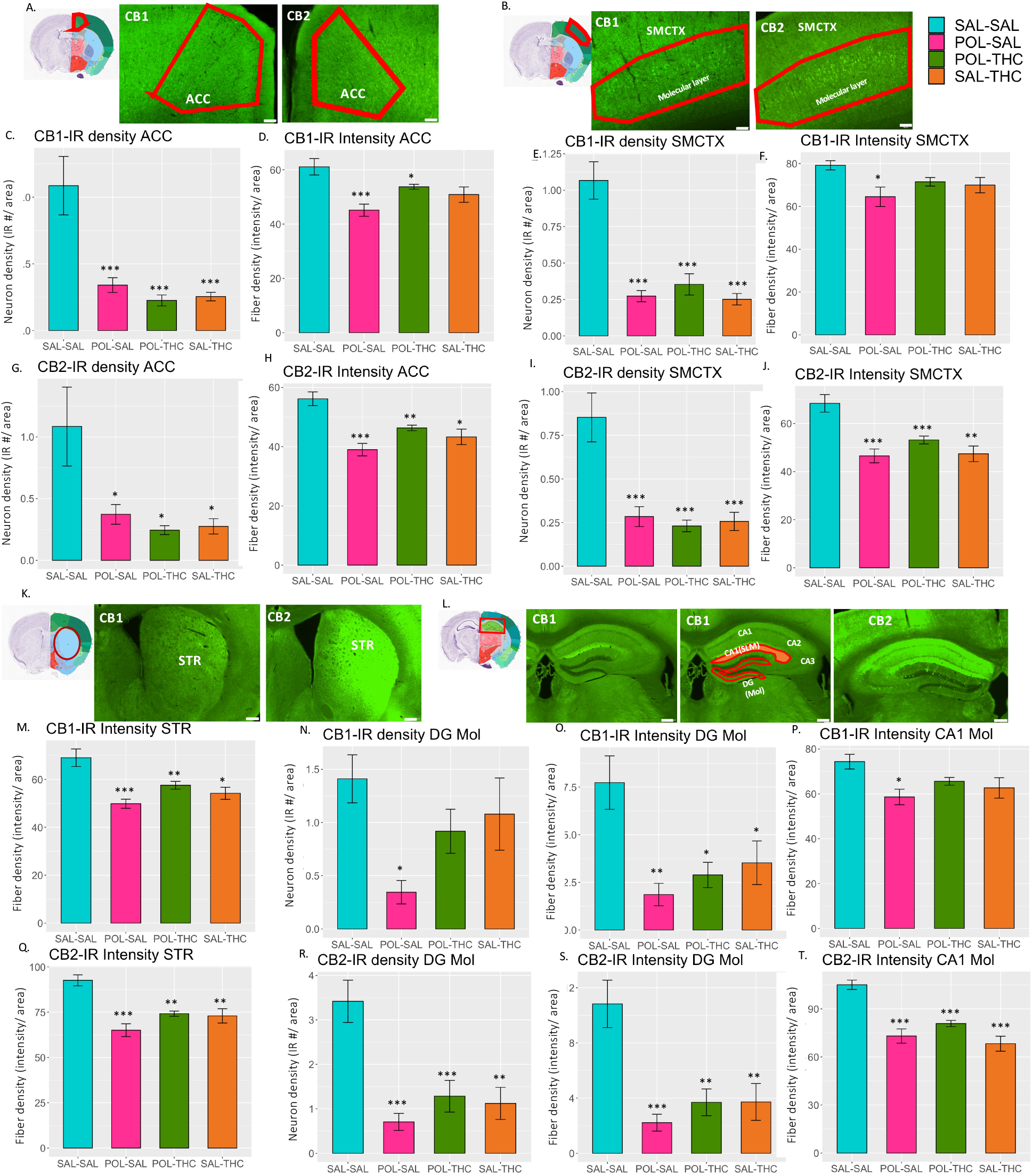
Decreases in the neuron cell and axonal fiber density of cells expressing CB1 and CB2 receptors due to risk factor exposure. **A**. The reference slice from the Allen Brain mouse reference atlas (https://mouse.brain-map.org/static/atlas), plus representative images of CB1 and CB2 stains are presented for the anterior cingulate cortex (ACC), with neuron cell density (**C**) and fiber density (**D**) for CB1 expression and CB2 expression (**G** neuron cell density and **H** fiber density). The same pattern is used for the somatosensory cortex (SMCTX; **B**) with CB1 and CB2 neuron cell density and fiber density (**EF** CB1, **IJ** CB2). For the striatum (STR; **K**) CB1 and CB2 fiber densities are represented in **M** and **Q**, respectively. Finally, for the hippocampus (**L**), results for the dentate gyrus molecular layer CB1 neuron cell density (**N**) and fiber density (**O**) are available for CB1 (**NO**) and CB2 (**RS**), while results for the CA1 molecular layer CB1 and CB2 fiber density are represented in **P** and **T**, respectively. Scale bars represent 100 μm for all regions except the hippocampus, where they represent 200 μm. *p<0.05, **p<0.001, ***p<0.0001 for Tukey’s HSD.

In the DG molecular layer, a subthreshold effect for CB1-IR (F(5,40)= 4.027, p=0.052) and significant effect for CB2-IR (F(5,40)=17.807, p=0.000136) cell density was observed. Tukey’s post hoc pairwise test revealed a significant decrease for POL-SAL relative to SAL-SAL for CB1-IR density, (p=0.0162), and a significant decrease in CB2-IR density for all 3 risk factor groups relative to SAL-SAL (p<0.001) (**Figure 5**). In these regions, similar patterns were observed for fiber density measures (presented in **Supplement 2.8**). Similarly, there was a significant interaction in CA1 molecular layer of the hippocampus for fiber density (cell density not quantified) for CB1-IR (F(5,40)= 6.184, p=0.0172) and CB2-IR (F(5,40)= 6.191, p=2.55e-07). CB1-IR fiber density was significantly reduced in POL-SAL relative to SAL-SAL (p<0.05), while all three risk factor groups had reduced CB2 fiber density relative to SAL-SAL (p<0.00001) (**Figure 5**). Summary statistics available in **Supplementary tables 5-20**.

## 4. Discussion

Converging lines of evidence across epidemiological and experimental studies have identified prenatal MIA-exposure and adolescent THC exposure as risk factors for neuropsychiatric illness. However, most individuals exposed to either risk factor alone do not develop psychiatric or neurodevelopmental disorders, which suggests that multiple exposures within an individual are required for disease onset (4,42). The work presented here is aimed at experimentally testing the multiple exposure hypothesis, by exposing MIA-primed offspring to a second hit: chronic adolescent THC exposure. Our results suggest that exposure to either prenatal MIA or adolescent THC were not sufficient to induce enduring neuroanatomical or behavioural alterations in adulthood, while combined exposure was, in some regions but not others. Behaviourally, only subtle, subthreshold alterations in anxiety-like and sensorimotor gating behaviours were observed in the group exposed to both risk factors. Interestingly, sex-differences were observed in patterns of brain-behaviour covariation, both for adult brain volume and for the within-subject brain volume change from post-treatment to adulthood, and adult behaviour. In several brain regions showing susceptibility to risk factor exposure (anterior cingulate and somatosensory cortices, striatum, hippocampus), striking decreases in the expression of CB1- and CB2-IR cells were observed in offspring exposed to either prenatal MIA, adolescent THC, or both, indicating that the effects of these risk factors may converge on the endocannabinoid system, amongst others. Our results show that a single risk factor may not be sufficient to cause enduring changes in gross neuroanatomy and behaviour, but that a combined exposure may be. Further, a single risk factor may affect the brain at finer scales of resolution.

Preclinical investigation of MIA-exposure has been critical to our understanding of neurodevelopmental disorders (6,43,44) identifying behavioural, neuroanatomical, and transcriptional alterations relevant to ASD and schizophrenia (44–49). Mice prenatally exposed to MIA had no statistically significant deviations in developmental trajectories, nor in anxiety-like, social, or sensorimotor gating behaviours. The lack of statistically significant effects was somewhat surprising, and not consistent with previous reports of significantly altered brain and behavioural development from our group (7) and others (37,49). However, exploration of the findings at an uncorrected (p<0.05) threshold identified altered trajectories in the striatum, hippocampus, lateral septum, and somatosensory cortex, wherein the MIA-exposed mice had an overshoot in the adolescent/early-adult period followed by a normalization in later adulthood (relative to SAL-SAL), in line with our previously published findings (7). The lack of behavioural alterations in adulthood is consistent with our previous findings; however, there are many reports of altered behaviour in adult MIA-exposed offspring across multiple domains (44). Some methodological differences may explain why we do not recapitulate our findings at the same level of statistical significance between this study and our previous one (7); most importantly, mice (lower n) were scanned only three times (compared to four in our previous work), which may have reduced our sensitivity to detect alterations in developmental trajectories (50). Additionally, the ages selected for MRI in this study did not perfectly match those of our previous work, which may have decreased our ability to capture the stages of greatest change across development. Significant brain structure alterations have been replicated by our group in adolescent/young adult (PND 35 and 60) MIA-exposed offspring indicating that we may have missed the vulnerable window in the current study (51). Finally, structural imaging was performed slightly differently between studies; here we used a cryogenically cooled surface coil with no contrast enhancement (manganese chloride administration 24 hours prior to scan), whereas in the previous study we used a quadrature volumetric coil, with contrast enhancement; perhaps differences in structural MRI acquisition could contribute to the lack of consistency in our findings.

Mice chronically exposed to THC in adolescence similarly had transient deviations in neurodevelopmental trajectories, with a peak difference in the adolescent/early-adult period. In various regions including the somatosensory cortex, medial preoptic area, globus pallidus, third ventricle, and reticular nucleus, THC exposed mice experienced a volume increase at the post-treatment timepoint (PND50), followed by a normalization at the later adult timepoint. In contrast, hippocampal volume decreased at the posttreatment timepoint, but also normalized. These results indicate that the chronic THC treatment may have induced neuroanatomical remodeling, but that given a few weeks without THC exposure, the brain normalizes relative to the control (SAL-SAL) exposed offspring. In agreement with the lack of neuroanatomical differences by the adult timepoint (PND85), we did not detect any behavioural alterations; had we tested these mice earlier in development or across other behavioural domains, we may have detected differences.

To our knowledge, no longitudinal rodent MRI studies exist examining the effects of chronic adolescent THC exposure on brain structure. Studies using other modalities do report neuroanatomical, neurochemical, and behavioural abnormalities relevant to the psychosis spectrum (36,38,52). Positron emission tomography studies of rats exposed to chronic THC report increased D2-like receptor availability in the dorsal striatum (53). Furthermore, chronic THC exposure in adolescence has been associated with decreased synaptic arborization in the rat hippocampus (36), and the prefrontal cortex (38). Impairments in cognitive flexibility, sensorimotor gating, and memory have all been identified in rodents chronically exposed to THC in adolescence (54–56).

Greater alterations to neurodevelopmental trajectories were observed in offspring exposed to both prenatal MIA and chronic adolescent THC. In the nucleus accumbens, anterior cingulate cortex, and entorhinal cortex, significant deviations in trajectory were observed, defined by enlarged volume in the POL-THC group relative to SAL-SAL at the post-treatment (PND50) scan. Similarly, to the SAL-THC group, these brain regions normalized in the absence of continued THC exposure by later adulthood (PND85). However, in the striatum and somatosensory cortex, the volume increase observed following the post-treatment scan was sustained into later adulthood, indicative of a more lasting neuroanatomical change. Similarly, in the subiculum, a volume decrease was observed following treatment, which was sustained into later adulthood. These regions may be more sensitive to the combined risk factor exposure; interestingly, they have been previously implicated in neurodevelopmental and neuropsychiatric disorders (57–62). Finally, investigation of the effects of THC in addition to MIA (POL-THC vs POL-SAL groups) revealed that mice exposed to both risk factors experience sustained growth of the bed nucleus of the stria terminalis, ventromedial thalamus, reticular nucleus and CA1, relative to mice exposed to MIA and adolescent SAL.

Behaviourally, we only observed subthreshold decreases in anxiety-like behaviours in our combined POL-THC group relative to SAL-SAL, potentially reflective of increased risk-taking behaviours. A subthreshold difference was also observed in sensorimotor gating behaviour, wherein the POL-THC mice performed worse at lower prepulse tones, but better at louder tones. Thus, although the combined risk factors were sufficient to induce neuroanatomical changes, they may not be sufficient to affect the behaviours we tested.

Strikingly, we observed dramatic decreases in CB-1 and CB-2 cell and fiber density in our regions of interest. Most pronounced were the decreases in the somatosensory cortex and ACC for both CB1 and 2. The somatosensory cortex showed consistent neuroanatomical alteration across the three risk factor groups, while the ACC showed sensitivity to both POL and POL-THC exposure. We observed a similar decrease in the fiber density of both CB1 and 2 staining in the striatum, another region affected by POL and POL-THC exposure. Finally, in the hippocampus we observed the most striking decreases in both CB1 and 2 cell and fiber density in the mice prenatally exposed to POL, while those exposed to adolescent THC (either prenatal SAL or POL) also had decreased density relative to SAL-SAL, but higher density relative to POL-SAL. Decreased CB1 receptor expression has been previously reported in the rodent brain (hippocampus and striatum) following both acute (63) and chronic THC exposure (64). Decreased sensitivity has also been observed in a number of rodent brain areas (cortex, hippocampus, and periaqueductal gray) following chronic use (31), as well as in chronic human users (temporal lobe, cingulate cortex, and nucleus accumbens (65). In contrast, increased expression has also been reported following short exposure to THC (66), indicating that the duration of exposure may modulate the directionality of changes to CB1 expression. Interestingly, previous work using PET imaging to longitudinally asses CB1 receptor expression in the offspring of MIA-exposed offspring (GD 15) found that MIA-exposed rats had lower CB1 receptor standard uptake values in the globus pallidus in adolescence, and lower values in the sensory cortex and hypothalamus in adulthood, relative to controls; these findings indicate that MIA may alter the cannabinoid system in adolescence and adulthood, potentially leading to greater sensitivity to drug use in this period (67).

The endocannabinoid system plays a crucial role in early brain development (68), and undergoes dynamic changes during the adolescent period, indicating that these two developmental windows may be particularly vulnerable to risk factors affecting this system (69). THC activates the endocannabinoid system precisely through the CB1 and CB2 receptors. The CB1 receptors are expressed in high concentrations in the hippocampus, amygdala, basal ganglia, many regions of the cerebral cortex and cerebellum (70,71). Activation of presynaptic CB1 receptors (via THC or endogenous endocannabinoids) suppresses GABA release (72), which can disrupt the development of pyramidal and parvalbumin-containing basket cell circuitry, creating an imbalance in excitatory-inhibitory signaling in the brain. Similar imbalances are thought to underlie psychosis (73), and have been identified in MIA-exposed offspring (74). Cannabis has also been shown to increase dopamine release acutely, which may underlie positive psychotic symptoms (hallucinations, delusions) (8). CB-1 receptors may also play a critical role in synaptic plasticity (75), and neuroinflammation, as CB1 receptors have been identified at low levels within microglial populations (76), and have been observed to affect glial cell function in response to cellular injury (77). CB2 receptors are highly expressed in the peripheral immune system and thought to have potent anti-inflammatory effects, making them interesting candidates for managing neuroinflammation as well (78). More recently, CB2 receptor expression has been identified in the cerebral cortex, hippocampus, striatum, amygdala, thalamic nuclei, periaqueductal grey, cerebellum and several brain stem nuclei of the rodent brain (40,78). These receptors may also play a functionally relevant role in the central nervous system, primarily through microglia (76). These may be interesting avenues for further investigation, as microglial activation has been implicated in many of the downstream neurodevelopmental aberrations due to MIA-exposure.

Few studies exist investigating the combined effects of the two risk factors studied here on offspring development. Previous work has identified decreased CB1 receptor availability due to prenatal MIA-exposure and adolescent cannabinoid exposure in the hypothalamus and sensory cortex, which are regions where we detect sustained alterations due to MIA and THC (79). Alterations to small non-coding microRNAs in the entorhinal cortex, associated with neurotransmission, cellular signaling, and schizophrenia, have been identified in rats exposed to both risk factors (80,81). Finally, alterations to the serotonergic (15) and dopaminergic systems (16) have both been identified in rats exposed to both risk factors; these neurotransmitter systems are relevant to the disease pathology of a number of neuropsychiatric disorders (82,83). These findings suggest that pathological changes may be occurring in response to the combined risk factors in some of the regions where we observe volume alterations. MIA-exposure has been shown to potentiate the effects of other environmental risk factors such as adolescent stress (84) or exposure to drugs of abuse (15), as well as genetic risk factors such as *DISC1* (85,86), *NRG1* (87), *NR4A2*, *TSC2* (88) causing greater deficits than either exposure alone (5,6).

Finally, our investigation of brain-behaviour covariation identified some underlying sex differences, not clearly uncovered in our univariate analyses. The pattern identifying in adulthood differentiated the male SAL-SAL offspring from all other male offspring exposed to either a single or a combined risk factor, whereas it did not differentiate the female offspring. This was defined by increased cortical, thalamic and cerebellar volume associated with decreased social interactions and sensorimotor gating. The assessment of within subject volume change post-treatment to adulthood with behaviour identified a similar behavioural pattern, associated with the somatosensory cortex striatum, hippocampus, and cerebellum; in this case, male mice were more similar in their expression of the pattern, whereas female SAL-THC offspring different from the other three groups. Overall, this analysis suggests that there may be interesting sex differences underlying the neurodevelopmental abnormalities associated MIA and THC exposure. This dimensional approach may parse variability better than categorical approaches (89) and has proven successful in our previous work (7,17,28)

Sex differences in response to MIA-exposure have been identified in human birth cohorts, showing that males are more likely to develop neurodevelopmental disorders following MIA-exposure to bacterial infection (90). Rodent studies also provide evidence for increased susceptibility in males, with reports of earlier deviations in neuroanatomy and behaviour based on longitudinal studies (49), as well as male-biased deficits in spatial working memory, PPI, and locomotion (91). Finally, altered synaptic function in the hippocampus and abnormal microglia activation have been reported in male MIA-exposed offspring, at a greater level than in females (30,92). Sex-biases have also been observed in response to cannabinoids; some evidence from human studies suggest that males may be more susceptible to the effects of cannabis use if they carry genetic risk for schizophrenia (12), and are also more likely to initiate use earlier than females (93). Rodent studies suggest that females may be more likely to develop behavioural despair, anhedonia, and catalepsy in response to THC treatment (94,95); these behavioural differences may, in part, be explained by differences in THC metabolism (96). Higher CB1 receptor density in males relative to females has been reported in regions such as the striatum, limbic system, and pituitary (97). Underlying sex-differences in several neurobiological systems including microglia and synaptic function associated with MIA-exposures, as well as CB1 receptor density and THC metabolism, may result in the sex-differences in the brain-behaviour alterations we observe.

Based on our multivariate results, we believe that future studies should be performed to specifically investigate sex differences in response to both risk factors, ensuring sufficient power to detect these complex statistical interactions. Further, acquiring MRI and behavioural data more frequently throughout the development of the offspring may allow us to fully investigate subtle changes. This may, in part, explain why we did not detect significant alteration in the MIA-exposed mice, and only detected subtle changes due to THC exposure. Furthermore, performing behavioural assays more proximal to the chronic THC exposure would allow us to uncover more acute behavioral alteration in response to treatment.

We find that exposure to a single risk factor, either prenatal MIA-exposure or chronic adolescent THC exposure are not sufficient to induce lasting neuroanatomical or behavioural changes in adulthood, although transient alterations were observed in adolescent/early-adult neuroanatomy. In contrast, exposure to both risk factors may induce lasting neuroanatomical deviations in some regions, but not others, while only inducing subthreshold behavioural alterations. Interestingly, exposure to either prenatal MIA or adolescent THC, or both, significantly decreases the expression of CB1 and 2 receptors in the brain. This points to the central role of the endocannabinoid system in both prenatal and adolescent brain development, as well as its sensitivity to disruptions due to risk factor exposure. In conclusion, our findings show that exposure to multiple risk factors may have greater effects on offspring brain volume throughout development than exposure to a single risk factor, but that CB1 and 2 receptor density is similarly decreased following exposure to a single or multiple risk factors. Finally, exposure to either one or two risk factors did not alter adult behaviour. These findings may further our understanding of how exposure to risk factors throughout development could increase the likelihood of developing neuropsychiatric illnesses.

## Acknowledgements

The authors would like to thank Dr. Joseph Rochford for sharing his expertise in data collection and analysis of the prepulse inhibition behaivioural task, and Dr’s Bruno Giros and Salah El Mestikawy for lending us their centrifuge. Data will be freely available for download on the zenodo platform (10.5281/zenodo.6373191). The authors would like to acknowledge their funding bodies, including the Canadian Institute of Health Research and Healthy Brains for Healthy Lives for providing support for this research. Additionally, we would like to thank the Fonds de Recherche du Québec en Santé for providing salary support for EG, KP, and MMC, as well as the Kappa Kappa Gamma Foundation of Canada for supporting EG’s salary.

