## Supplementary materials and methods for "Investigating the “two-hit hypothesis”: effects of prenatal maternal immune activation and adolescent cannabis use on neurodevelopment in mice"

### 1. Methods

#### 1.1. Animals

C57BL/6J mice were bred in our facility under a 12 hour light cycle (8 am-8 pm), with food and water access *ad libitum*. Females and males of breeding age (8-12 weeks) were placed in new cages (1:1 ratio) for up to 2 days until seminal plug was observed. This was considered gestational day (GD) 0. Each female was weighed and moved to a new cage. Animals were weighed again on injection day to confirm pregnancy. Weights were recorded on scan days (PND 25, 50, 85) and during chronic treatment (PND 28-45).

#### 1.2 Assessment of maternal cytokines levels

In a separate group of dams, poly I:C or saline was injected as described above (n=4 GD9-POL, n=5 GD9-SAL). Three hours following injection, dams were sacrificed by decapitation without euthanasia, and trunk blood was collected in a 1.5mL Eppendorf tube. The blood was allowed to coagulate at room temperature for 30 minutes, and was then centrifuged for 10 minutes at 4℃, with 2000 revolutions per minute. Serum was collected and stored at -80℃ until ready for analysis. Serum samples were shipped to the University of Maryland Core Cytokine Facility (http://www.cytokines.com/) for multiplex ELISA to measure levels of IL-6, TNF-alpha, IL-1**β**, IL-10. We chose to use a separate group of dams to ensure we could collect enough blood for analysis, and so as not to introduce an additional stressful experience for the dam, thereby potentially confounding the neurodevelopmental trajectory of offspring. Detection ranges were as follows IL-6 (1.95-8000 pg/ml), TNF-alpha (0.85-3500 pg/ml), IL-1**β** (3.75-15000 pg/ml), IL-10 (5-20000 pg/ml).

#### 1.3 Validation of THC solution

A stock solution of 2 g of pure delta-9-THC 10 mg/ml in ethanol was obtained (Cayman Chemicals, Ann Arbor, MI, USA). To prepare a physiologically compatible solution, vacuum drying was used to extract the THC from ethanol and prevent conformational changes to the compound. Pure THC was then dissolved in a 1:18 cremophor:saline solution, resulting in a 1:1:18 THC:cremophor:saline solution (the lipophilic cremophor ensures the THC remains in solution).

Plasma levels of THC were measured in a separate cohort of mice with gas-chromatography mass spectrometry (GCMS) (1) coupled to a flame ionization detector. Adult C57BL/6J mice were injected with dosages of either 2.5 mg/kg (N=5, 2F/3M), 5 mg/kg (N=5, 2F/3M), 10 mg/kg (N = 5, 3F/2M), or cremophor:saline control (N = 5, 3F/2M). One hour post injection, mice were euthanized by live decapitation and trunk blood was collected for analysis in 1.5 mL EDTA Eppendorf tubes containing chilled EDTA (final concentration of 5mM). Samples were centrifuged for 15 minutes (1500 rotations) at 4℃. Plasma was collected in separated Eppendorf tubes and stored at -80℃ until analysis. 1 mg/mL THC in methanol was purchased from Millipore Sigma-Canada (Product Number T4764) to serve as the standard for the GCMS. Samples were tested for the presence of delta-9-THC, as well as two THC metabolites, 11-hydroxy-delta-9-THC and 11nor-9carboxy delta-9-THC. Results indicated for all dosages THC and its metabolites were present at values within the expected bounds of error across the groups (**Supplementary Table 1**).

### 1.4 Magnetic resonance imaging

Structural images (n=245) were exported as DICOM, converted to MINC format, preprocessed, and visually inspected by two independent raters for quality control (QC). Three scans from one subject were excluded due to hydrocephalus and one scan was excluded due to motion (n=242). All scans per subject were registered to create a subject average (first-level). These were registered using a group-wise averaging technique to create a study average (second-level). The final average provides voxel correspondence between subjects, which allows for comparison of local individual changes across subjects (2). This was done using the antsMultivariateTemplateConstruction2.sh tool (https://github.com/CoBrALab/twolevel_ants_dbm)(3). Voxel-level volume changes were captured in the Jacobian determinant of the deformation field derived from the image registration. Both relative and absolute Jacobian determinants of the deformations fields of the first level were resampled into the second level average space to perform statistics (4). Relative Jacobians explicitly model only the non-linear part of the deformations to remove residual global linear transformation attributable to differences in total brain size and were used for subsequent statistical analysis. Absolute Jacobians include overall linear transformations. Jacobian determinants (5) of the first-level deformation fields were resampled into final average space and blurred (0.2 mm Gaussian 3D kernel) prior to statistical analyses (6,7). All trajectory plots (**Figure 2**) were created using the Effect function in R to correctly reflect the statistical model, accounting for covariates (https://github.com/CoBrALab/documentation/wiki/Properly-plotting-a-lm-or-lmer-model-predicted-curve-in-R-with-ggplot).

### 1.5 Behavioural tests

#### 1.5.1 Open field test

This test was used to assess exploratory and anxiety-like behaviours. Mice were gently placed in the center of a 45 x 45 cm rectangular light grey arena and allowed to explore for 15 minutes, while being video recorded. Distance traveled and time spent in conceptually partitioned arenas was measured: a center zone (40% of the total area), and the perimeter (corners and edges). Distance traveled in the center zone relative to total distance traveled was used as the main assay, as was cumulative duration spent in the center zone.

#### 1.5.2 Three chambered social preference and social novelty task

Mice were habituated (10 minutes) under red light to a three-chamber plastic box (26 (l) x 21.6 (w) x 21.6 (h) cm) with divider panels that have open doors, with a wire container (9.5 (h) 7.6

(d) cm) in each of the two extreme chambers. To measure social preference (10 minutes), time spent interacting with a stranger mouse was compared to that with a non social object using the following social preference index formula 1:

```
([time spent sniffing intruder 1 zone] / [time spent sniffing
object zone + time spent sniffing intruder 1 zone]) - 0.5. (1)
```

Similarly, to measure social novelty (10 minutes), the nonsocial object was replaced with another stranger mouse, and a social novelty index was calculated as formula 2:

```
([time spent sniffing intruder 2 zone] / [time spent sniffing
intruder 1 zone + time spent sniffing intruder 2 zone]) - 0.5
(2)
```

Stranger mice were the same strain, sex, and similar age (within 2 weeks) of the test mice, and were habituated to the wire containers (20 minutes twice a day) for two days prior to the test.

#### 1.5.3 Prepulse inhibition

Prepulse inhibition (PPI) to acoustic startle was measured using commercially available startle chambers (San Diego Instruments, San Diego, CA) consisting of a Plexiglass chamber (8 cm diameter, 16 cm long) mounted on a Plexiglass base with a sound-attenuating chamber, and a speaker located in the ceiling of the chamber (24 cm above the animal) to provide the background noise (70dB) and the acoustic stimuli. A piezoelectric accelerometer fixed to the animal enclosure frame was used to detect and transduce motion resulting from the animal's startle response. A microcomputer using a commercial software package by SR-LAB was used to control pulse parameters, and digitize (0-4095), rectify, and record the stabilimeter readings. Animals were placed in the Plexiglass restrainers, and after 5 minutes of acclimatization. Mice underwent a total of 50 trials (5-30 s intertrial duration).

Startle magnitude to a 50 ms 120 dB stimulus, in absence of prepulse, was measured in the first 8 and final 7 trials. For the middle 35 trials, the startle tone was either presented alone, or preceded by a 30ms prepulse stimulus ranging from 3-15 dB above background noise (73-85 dB) and varying randomly between trials in 3 dB increments (5 trials per prepulse stimulus). A measure of maximum and average startle response was derived from the 100 1 ms readings taken starting from the beginning of the startle stimulus onset. Percent PPI from each prepulse intensity formula 3:

```
(averaged over trials) was calculated using the formula:
[(startle response - prepulse response) / startle response] x
100 (3)
```

Differences in maximum startle amplitude were investigated.

### 1.6 Statistical modeling

#### 1.6.1 Weight data

To determine whether chronic THC exposure affected weight of the animals, we performed a linear-mixed effects model testing for a prenatal treatment-by-postnatal treatment-by-age interaction, with sex as a covariate, and mouse as a random intercept. We focused on the PND 25 - 50 ages to focus on the more acute effects of treatment

**Weight model:** $Y_{subject,j} = \boldsymbol{\beta}_0 + \boldsymbol{\beta}_1 sexF_{subject,j} + \boldsymbol{\beta}_2 prenatalPOL_{subject,j} + \boldsymbol{\beta}_4 PostnatalTHC_{subject,j} + \boldsymbol{\beta}_5 Age_{subject,j} + \boldsymbol{\beta}_6 prenatalPOL{:}postnatalTHC_{subject,j} + \boldsymbol{\beta}_7 prenatalPOL{:}Age_{subject,j} + \boldsymbol{\beta}_8 postnatalTHC{:}Age_{subject,j} + \boldsymbol{\beta}_9 Age{:}prentalPOL_{subject,j}{:}postnatalTHC_{subject,j} + \mathbf{b_1} subject + \mathbf{b_2} litter + \epsilon_{subject,j}$

Y= outcome measures (i.e. weight in g); $\boldsymbol{\beta}_i$= fixed effect coefficient; $\boldsymbol{\beta}_0$ = equation intercept; **b** = random predictor; **ϵ** = random error; **j** = repeated measure per subject; **:** = interaction; prenatalPOL= early polyI:C group relative to SAL as reference; postantalTHC = adolescent THC group relative to SAL as reference; SexF = female sex relative to male as reference

#### 1.6.2 Neuroimaging data

A voxel-wise linear mixed-effects model was run at every voxel in the brain with the following two models, the first for assessing overall group-by-age interactions, and the second for investigating for sex differences:

**MIA model:** $Y_{subject,j} = \boldsymbol{\beta}_0 + \boldsymbol{\beta}_1 sexF_{subject,j} + \boldsymbol{\beta}_2 groupPOL_{subject,j} + \boldsymbol{\beta}_4 age(ns,2)1_{subject,j} + \boldsymbol{\beta}_5 age(ns,2)2_{subject,j} + \boldsymbol{\beta}_6 ns(age,2)1{:}groupPOL_{subject,j} + \boldsymbol{\beta}_7 ns(age,2)2{:}groupPOL_{subject,j} + \mathbf{b_1} subject + \mathbf{b_2} litter + \epsilon_{subject,j}$

**THC model:** $Y_{subject,j} = \boldsymbol{\beta}_0 + \boldsymbol{\beta}_1 sexF_{subject,j} + \boldsymbol{\beta}_2 groupTHC_{subject,j} + \boldsymbol{\beta}_4 age(ns,2)1_{subject,j} + \boldsymbol{\beta}_5 age(ns,2)2_{subject,j} + \boldsymbol{\beta}_6 ns(age,2)1{:}groupTHC_{subject,j} + \boldsymbol{\beta}_7 ns(age,2)2{:}groupTHC_{subject,j} + + \mathbf{b_1} subject + \mathbf{b_2} litter + \epsilon_{subject,j}$

**POL THC model:** $Y_{subject,j} = \boldsymbol{\beta}_0 + \boldsymbol{\beta}_1 sexF_{subject,j} + \boldsymbol{\beta}_2 groupPOL\text{-}THC_{subject,j} + \boldsymbol{\beta}_4 age(ns,2)1_{subject,j} + \boldsymbol{\beta}_5 age(ns,2)2_{subject,j} + \boldsymbol{\beta}_6 ns(age,2)1{:}groupPOL\text{-}THC_{subject,j} + \boldsymbol{\beta}_7 ns(age,2)2{:}groupPOL\text{-}THC_{subject,j} + + \mathbf{b_1} subject + \mathbf{b_2} litter + \epsilon_{subject,j}$

**Sex differences:**

**MIA model:** $Y_{subject,j} = \boldsymbol{\beta}_0 + \boldsymbol{\beta}_1 sexF_{subject,j} + \boldsymbol{\beta}_2 groupPOL_{subject,j} + \boldsymbol{\beta}_4 age(ns,2)1_{subject,j} + \boldsymbol{\beta}_5 age(ns,2)2_{subject,j} + \boldsymbol{\beta}_6 ns(age,2)1{:}groupPOL_{subject,j} + \boldsymbol{\beta}_8 ns(age,2)2{:}groupPOL_{subject,j} + \boldsymbol{\beta}_9 ns(age,2)1{:}groupPOL_{subject,}{:}sexF_{subject,jj} + \boldsymbol{\beta}_{10} ns(age,2)2{:}groupPOL_{subject,j}{:}sexF_{subject,j} + \mathbf{b_1} subject + \mathbf{b_2} litter + \epsilon_{subject,j}$

**THC model:** $Y_{subject,j}$ = $\boldsymbol{\beta}_0$ + $\boldsymbol{\beta}_1$sexF$_{subject,j}$ + $\boldsymbol{\beta}_2$groupTHC$_{subject,j}$+ $\boldsymbol{\beta}_4$age(ns,2)1$_{subject,j}$ + $\boldsymbol{\beta}_5$age(ns,2)2$_{subject,j}$ + $\boldsymbol{\beta}_6$ns(age,2)1:groupTHC$_{subject,j}$ + $\boldsymbol{\beta}_7$ns(age,2)2:groupTHC$_{subject,j}$ + $\boldsymbol{\beta}_8$ns(age,2)1:groupTHC$_{subject,}$:sexF$_{subject,jj}$ + $\boldsymbol{\beta}_9$ns(age,2)2:groupTHC$_{subject,j}$:sexF$_{subject,j}$ + $\mathbf{b}_1$subject + $\mathbf{b}_2$litter + $\epsilon_{subject,j}$

**POL THC model:** $Y_{subject,j}$ = $\boldsymbol{\beta}_0$ + $\boldsymbol{\beta}_1$sexF$_{subject,j}$ + $\boldsymbol{\beta}_2$groupPOL-THC$_{subject,j}$+ $\boldsymbol{\beta}_4$age(ns,2)1$_{subject,j}$ + $\boldsymbol{\beta}_5$age(ns,2)2$_{subject,j}$ + $\boldsymbol{\beta}_6$ns(age,2)1:sexF$_{subject,j}$ + $\boldsymbol{\beta}_7$ns(age,2)2:sexF$_{subject,j}$ + $\boldsymbol{\beta}_8$ns(age,2)1:groupPOL-THC$_{subject,j}$ + $\boldsymbol{\beta}_9$ns(age,2)2:groupPOL-THC$_{subject,j}$ + $\boldsymbol{\beta}_{10}$ns(age,2)1:groupPOL-THC$_{subject,j}$:sexF$_{subject,j}$ + $\boldsymbol{\beta}_{11}$ns(age,2)2:groupPOL-THC$_{subject,j}$:sexF$_{subject,j}$ + $\mathbf{b}_1$subject + $\mathbf{b}_2$litter + $\epsilon_{subject,j}$

Y= outcome measures (i.e., blurred absolute Jacobian determinants); $\boldsymbol{\beta}_i$= fixed effect coefficient; $\boldsymbol{\beta}_0$ = equation intercept; **b** = random predictor; **ϵ** = random error; **j** = repeated measure per subject; **:** = interaction; POL= early polyI:C group relative to SAL-SAL as reference; THC = adolescent THC group relative to SAL-SAL as reference (or POL SAL as reference); POL-THC = POL-THC group relative to SAL-SAL as reference; SexF = female sex relative to male as reference

### 1.6.3 Behavioral data

For PPI data across increasing decibel levels, differences in trajectory of sensorimotor gating changed over increasing prepulse level were modeled with a second order natural spline, both overall and for sex differences, as described above for the MRI data.

### 1.7 Partial least squares significance and reliability assessment

The behaviour matrix was z-scored and correlated to the brain matrix to create a brain-behaviour covariance matrix. Singular value decomposition was applied to brain-behaviour matrix to generate a set of orthogonal latent variables (LVs), which describe linked patterns of covariation between the input brain and behaviour matrices. Permutation testing and bootstrap resampling (n=1000 each) were used to assess LV significance and reliability. Outliers in brain-behaviour scores were determined by computing the outliers minimum (quartile 1 - 1.5x interquartile range) and maximum (quartile3 + 1.5x interquartile range) for each score (3 [2 male, 1 female] SAL-THC and 2 [1 male and 1 female] POL-SAL were removed).

***Permutation testing:*** was used to assess the statistical significance of each LV wherein the rows (subjects) of the brain data matrix were randomly shuffled to 1) nullify dependencies between brain and behaviour (n=1000 repetitions) and 2) generate a null distribution of possible brain-behaviour correlations. SVD was applied to these "null" correlations, generating a distribution of singular values under the null hypothesis. The probability that a permuted singular value exceeds the original, non-permuted singular value allows us to generate the p-value(8,9). A threshold of $p<0.05$ was used (95% or greater chance that the singular value of the non permuted data exceeds that of a permuted singular value).

***Bootstrap resampling:*** was applied to assess the contribution of individual brain and behaviour variables to each LV. Subjects (rows for both X and Y matrices) were randomly sampled and replaced (n=1000) to generate a set of resampled correlation matrices to which SVD was applied to generate a sampling distribution for each weight of the singular vectors. The ratio of each singular vector weight and its bootstrap-estimated standard error were used to calculate a "bootstrap ratio" for each voxel. Voxels that make large contributions to certain patterns can therefore be identified by large bootstrap ratios.

### 1.8 Immunohistochemistry

#### 1.8.1 Perfusion

Mice from four groups were perfused [SAL-SAL (n=5), POL-SAL (n=6), POL-THC (n=6) and SAL-THC (n=6)]. All mice were anaesthetized with a cocktail of ketamine/xylazine mixture and intracardially perfused with cold 1X phosphate buffered saline (PBS) and followed by 4% paraformaldehyde in 1X PBS. Whole brains were removed and post-fixed in the same fixative overnight. A small hole was made in the left hemisphere to differentiate left from right in subsequent analyses. Mouse brains were transferred to 1X PBS and kept at 4$^{o}$C for more than one year. One week before sectioning, all brains were transferred to 30% sucrose in 1X PBS for cryoprotection. At the same time, a marker to differentiate the left from the right side of the brain was made for each mouse brain. Embedded in M1 embedding medium (Fisher), brain blocks were cut into 50 µm-thick coronal sections on a cryostat (Leica). Sections were mounted on untreated clean glass slides (Fisher) and stored in -80$^{o}$C freezer until immunofluorescent staining.

#### 1.8.2. Immunofluorescent staining

On the day of free-floating immunofluorescent staining, brain sections were taken out of the freezer and brushed into the wells of a 24-well culture plate. Sections were incubated in 10% normal goat serum (NGS) diluted in 1X PBS containing 0.05% Triton-100 (PBS + T) for 2 hours. Then sections were incubated in polyclonal rabbit antiserum raised against cannabinol (CB)1 or CB2 receptors (1:250, Abcam) for 18 h at room temperature. Sections were next incubated in Alexa Fluor-488 conjugated goat anti-rabbit IgG (1:500, Invitrogen) in the darkness for 2 hours. Then sections were incubated in DAPI-PBS (0.5µg/ml) for 10 min in the darkness to stain the nuclei of cells. Between incubations, brain sections were washed thoroughly in PBS + T twice. Finally, sections were mounted on the pre-cleaned super plus glass slides (Fisher) and cover-slipped with anti-fading mounting medium 0.02% Miowol solution (Sigma) and observed under a fluorescent microscope (Olympus).

#### 1.8.3 Capture of CB1- and CB2-immunoreactive (IR) receptors images and quantification

The setup for image capturing was the same for all sections (~3 per mouse), analyzing left and right hemisphere for each mouse separately. For each brain region, images were captured with the same threshold of GFP illumination or exposure time for all sections across all groups on the fluorescent microscope. Axonal fibers expressing CB1- and CB2-immunoreactive (IR) receptors were distributed in the CA1-3 of hippocampus and striatum. The images of these CB1- and CB2-IR axonal terminals in these brain regions were captured at the 4X magnification. CB1- and CB2-IR neurons localized in the primary sensory cortex, anterior cingulate cortex, and in the molecular layer of dentate gyrus were captured at the 10X magnification.

The quantification of CB1- and CB2-IR varied by region, depending on what type of cell/process was CB1-positive. In the striatum (image 50 of the Allen Brain Atlas (ABA) mouse reference atlas; Bregma +0.38 mm) and molecular layer of CA1 of hippocampus (image 76; Bregma -2.80mm), average optical intensity of axon terminals was measured automatically using Fiji ImageJ (average values ranged from 1-256). The outlines of these brain regions were traced based on the ABA mouse reference atlas. For each mouse, mean average intensity was computed across all sections in the counting area, which gave us a measure of fiber density.

In the primary somatosensory cortex (image 62-64; Bregma -0.94 mm), anterior cingulate cortex (image 62-64; Bregma -0.94 mm) and the molecular layer of the dentate gyrus of the hippocampus (image 76; Bregma -2.80 mm), CB1- and CB2-IR neurons were counted directly using the cell counter Karma. Three traced areas in the molecular layers of the primary sensory cortex and in the anterior cingulate cortex were randomly selected; the area and signal intensity of these regions was measured automatically using Fiji's ImageJ. CB1- and CB2-IR neurons were counted manually only within the traced and measured areas. Only CB1- and CB2-IR neurons with a clear nucleus within the three measured areas were counted manually. Given the small size of the molecular layer of the dentate gyrus, the whole region was traced and measured using SigmaScan. In this region, CB1- and CB2-IR neurons were sparser, but very intensely immunostained, making nuclei less visible (relative to the bring processes). Thus, cells with clear neuron profiles, but with or without branches, were also counted. The ratio of the cell number vs. the measured area was calculated. The ratio of cells/area from the three measured regions of primary sensory cortex of either side were averaged. Then the ratio values from three brain sections of either side were also averaged and determined for each mouse brain. The mean neuron counts and the mean ratios of IR-neuron number vs. area were determined for each group.

### 2. Results

#### 2.1 Poly I:C injection increases pro-inflammatory cytokines in the dam

We observed an increase in levels of all three pro-inflammatory cytokines measured, IL-6, IL-1**β**, and TNF-ɑ, as well as in the anti-inflammatory cytokine IL-10 in a separate cohort of pregnant dams 3-hours post poly I:C injection on GD9 relative to saline control on GD9. Values are summarized below (Table 2).

**Supplementary Table 1.** Maternal serum cytokine levels for our 2 treatment groups, mean [range]

| | **IL-1β (pg/ml)** | **IL-6 (pg/ml)** | **IL-10 (pg/ml)** | **TNF-ɑ (pg/ml)** |
|---|---|---|---|---|
| **SAL** (n=5) | 21.578 [0.64-96.62] | 45.544 [30.58-56.59] | 21.862 [0.64-56.22] | 14.296 [0.64-26.05] |
| **POL** (n=4) | 36.3325 [0.62-142.81] | 5819.96 [4552.18-6512.84] | 135.845 [56.73-166.50] | 75.095 [45.72-90.51] |

#### 2.2 THC plasma concentrations in blood reflected injected dose

GCMS analysis of blood plasma THC concentrations confirmed that our doses were delivering a comparable amount in the mouse bloodstream 1-hour post-injection. Concentrations of other metabolites were below detection threshold (Table 3).

**Supplementary Table 2.** Mean plasma THC metabolite levels for satellite groups of animals, n/d = not detected

| | **d-9THC concentration (ng/ml)** | **11OH d-9THC (ng/ml)** | **11nor-9carboxy d-9THC (ng/ml)** |
|---|---|---|---|
| **10 mg/kg solution injection** | 9.6 mg/kg | n/d | n/d |
| **2.5 mg/kg solution injection** | 2.8 mg/kg | n/d | n/d |
| **5 mg/kg solution injection** | 5.8 mg/kg | n/d | n/d |

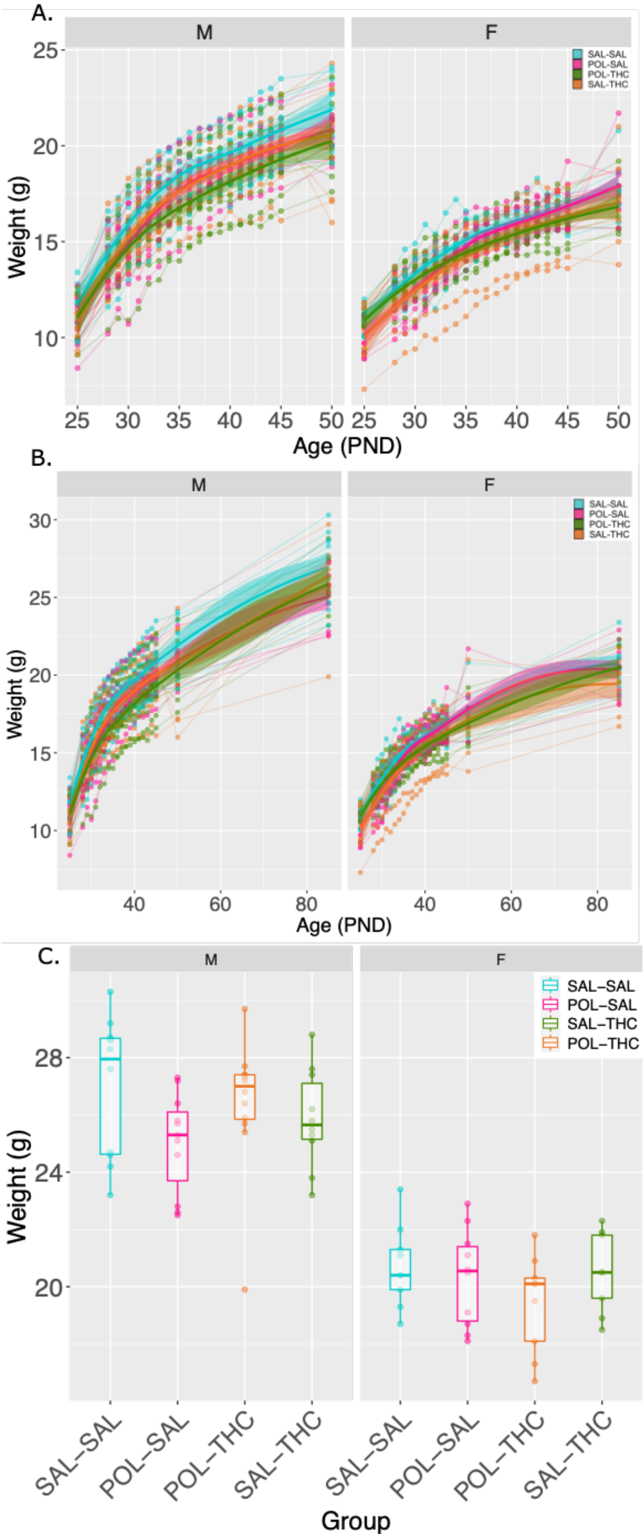

### 2.3 Weights

There was no significant three-way interaction between prenatal treatment, postnatal treatment and age. Significant two-way interactions for the effect of prenatal POL treatment and age (t=2.536, p=0.011) and postnatal THC treatment and age (t=-2.946, p=0.003) were observed, as were main effects for prenatal POL (t=-2.894, p=0.004), age (t=43.324 p=< 2e-16), and sex (t=-10.340, p=3.78e-16) (**Supplementary Figure 1**).

**Supplementary figure S1**. Group weights. **A.** Weights (g) for all 4 groups plotted from the pre- to post-treatment scan timepoint, with weight measures taken at PND 25, 28-45, and 50. **B**. Same data as in A, with the addition of weight measures taken at the final scan timepoint, PND 85. **C**. Weights for the PND 85 final scan timepoint alone.

### 2.4 Overview of neuroanatomical results

Pairwise comparisons for groups exposed to either one or both risk factors relative to control offspring revealed greater neuroanatomical differences in the combined risk factor group, POL THC (**Supplementary figure S2**).

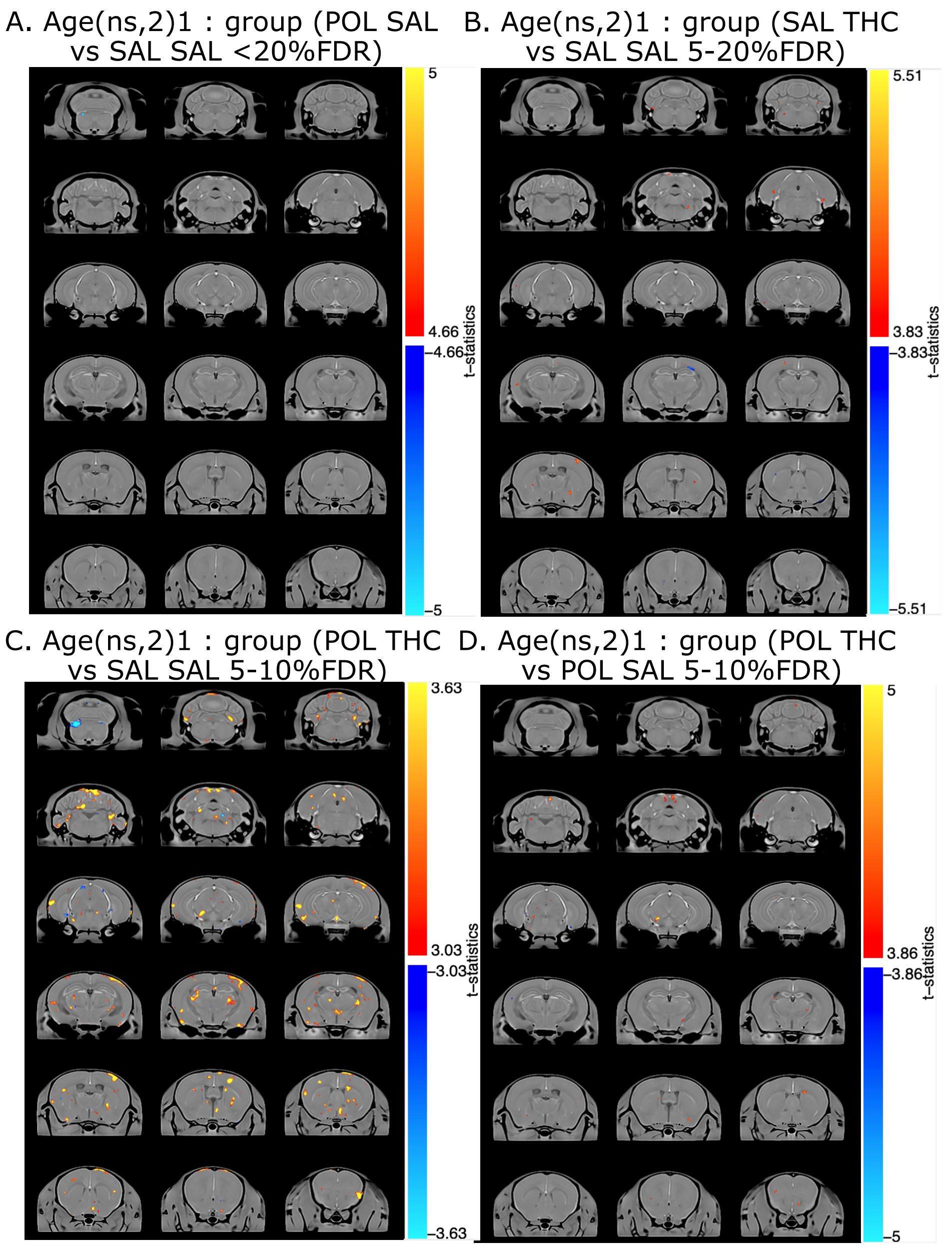

**Supplementary figure S2.** Overview of neuroanatomical changes in each group. T-statistic maps for the interaction between groups and the first order natural spline of age (*age(ns,2)1:group*) are displayed on the population average for comparison between POL-SAL and SAL-SAL (**A**), SAL-THC and SAL-SAL (**B**), POL THC and SAL-SAL (**C**), and POL-SAL and POL-THC (**D**). False discovery rate (FDR) correction thresholds highlighted for each panel.

### 2.5. Neuroanatomical alterations due to THC for POL offspring

For completeness, effects of the THC above and beyond those of MIA were explored by comparing offspring exposed both prenatally to POL and postnatally to SAL (POL-SAL), to those exposed prenatally to POL and postnatally to THC (POL-THC). A significant (*group:ns(age,2)1)* interaction (t=4.766, 10%FDR) was observed in the bed nucleus of the stria terminalis (BNST), the ventromedial thalamus, CA1 of the hippocampus, and the reticular nucleus. In all regions, the POL THC group started with a larger volume than POL SAL between PND 25 and 50, which decreased at PND 85. No sex differences were observed.

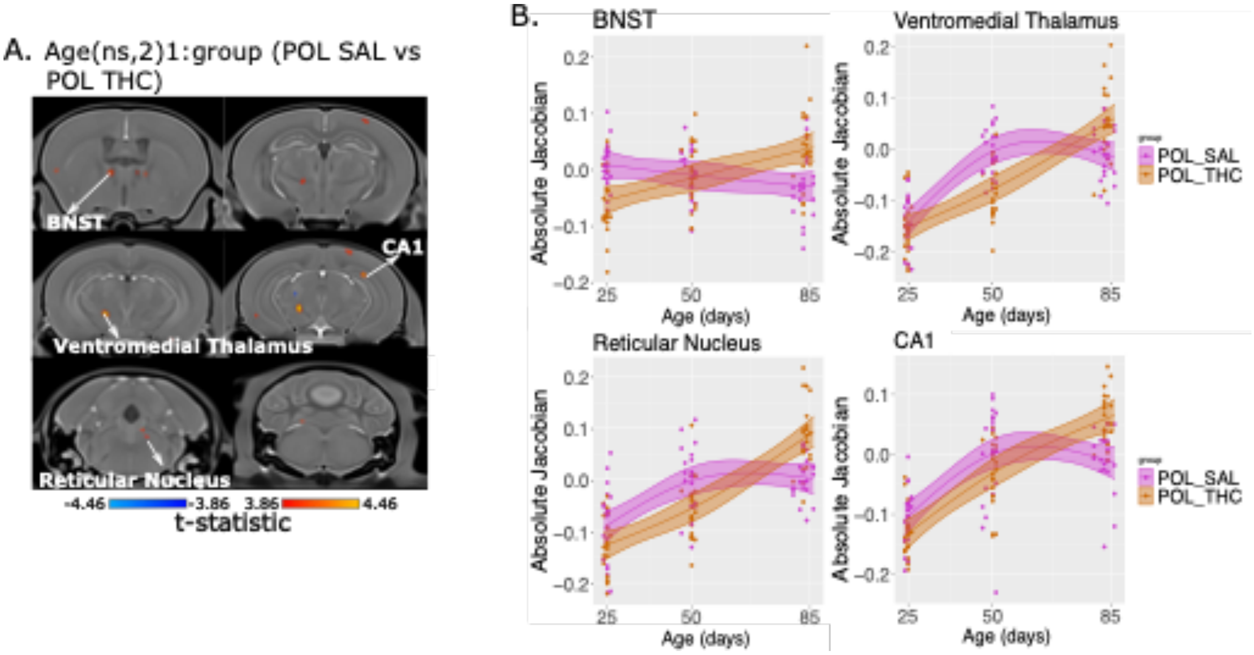

**Supplementary Figure S3.** Neuroanatomical alteration due to adolescent THC-exposure in MIA-exposed offspring exclusively. **A**. t-statistic map of group (POL THC vs POL SAL) by age (first order natural spline of age) thresholded between 5% FDR (top, t=4.46) and 10% FDR (bottom, t=3.86). **B**. Plot of peak voxels selected from regions of interest highlighted in **A**, wherein age is plotted on the x-axis, and the absolute Jacobian determinants plotted on the y-axis. Trajectories reflect the statistical model used (see **2.4.2**).

### 2.6 Behaviour

Overall, no statistically significant results were observed in our behavioural tests, however a few subthreshold effects were observed, as described in the main text (**3.4**) and in **supplementary table 4** below. No significant differences observed between POL THC and POL SAL ((**A**))

**Supplementary Table 3. Summary of all behavioural results for all group comparisons.** T-values, p-values (uncorrected) and q-values (corrected) are bolded if they survive Bonferroni correction (q-value=p<0.0125).

| | **POL-SAL vs. SAL-SAL** | **SAL-THC vs. SAL-SAL** | **POL-THC vs. SAL-SAL** | **POL-THC vs. POL-SAL** |
|---|---|---|---|---|
| **OFT** | t=0.943, p=0.361, q=1.000 | t=1.559, p=0.129, q=0.516 | t=2.344, p=0.041, q=0.164 | t=1.070, p=0.292, q=1.000 |
| **SOPT** | t=-0.496, p=0.632, q=1.000 | t=-1.217, p=0.233, q=0.932 | t=-0.977, p=0.335, q=1.000 | t=-0.345, p=0.732, q=1.000 |
| **SONT** | t=-1.990, p=0.054, q=0.216 | t=-0.017, p=0.9866, q=1.000 | t=-0.937, p=0.355, q=1.000 | t=0.733, p=0.468, q=1.000 |
| **PPI** | **Overall:** t=-0.148, p= 0.885, q=1.000 **By PP tone (*ns(level, 2)1:group)*** t=-0.560, p=0.577, q=1.000 | **Overall:** t=-0.013, p=0.990, q=1.000 **By PP tone (*ns(level, 2)1:group*)** t=1.268, p=0.208, q=0.832 | **Overall:** t=0.091, p=0.930, q=1.000 **By PP tone (*ns(level, 2)1:group*)** t=2.239, p=0.027, q=0.108 | **Overall:** t=-0.370, p=0.714, q=1.000 **By PP tone (*ns(level, 2)1:group*)** t=0.177, p=0.860, q=1.000 |

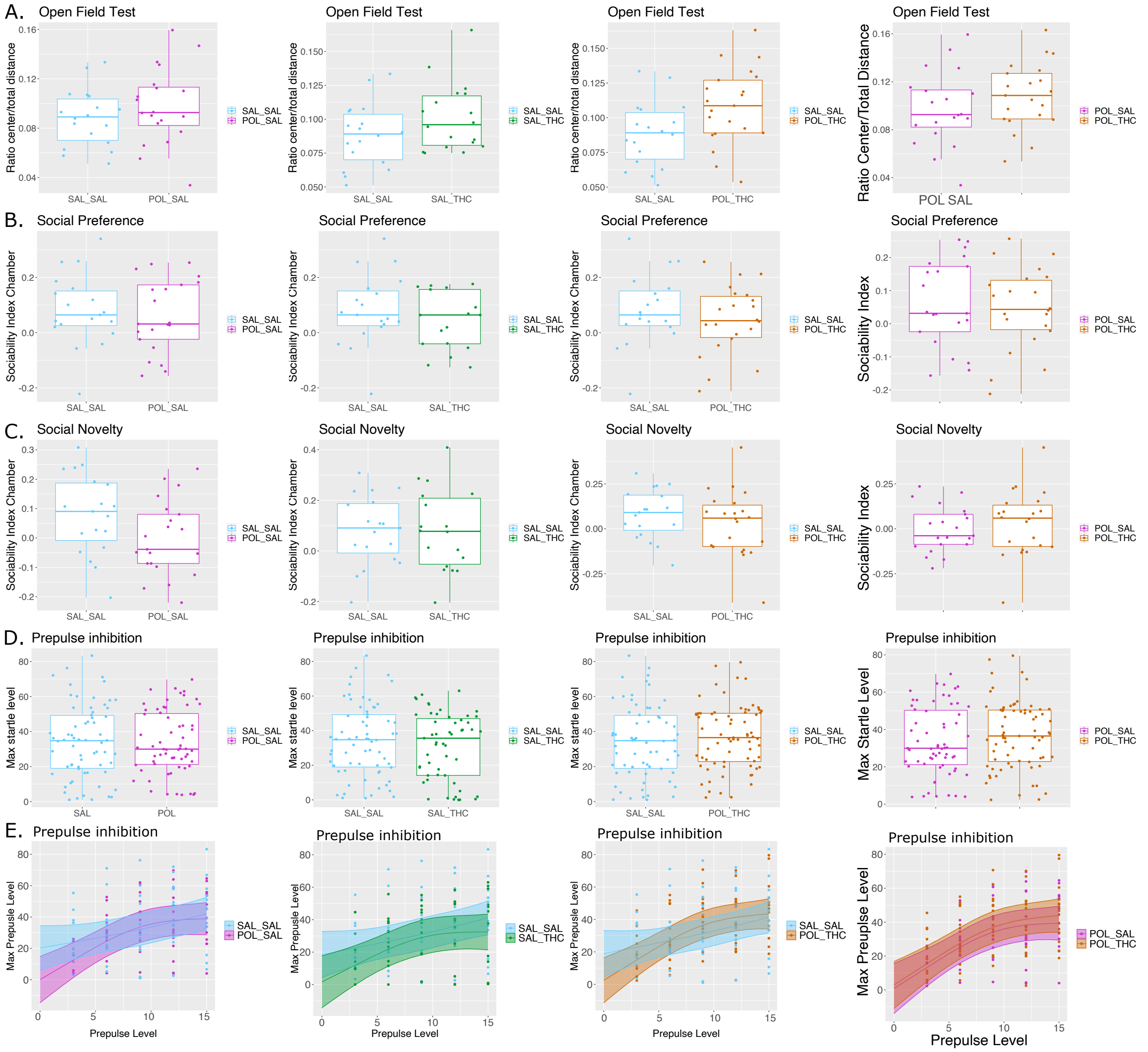

**Supplementary figure S4.** Prenatal MIA-exposure and/or adolescent THC exposure do not affect adult behaviour. Behavioural results for all treatment groups: SAL-SAL (cyan), POL-SAL (magenta), SAL-THC (green), POL-THC (orange). For all boxplots the midline represents the median of the data, the box represents the interquartile range, with whiskers denoting the full range of the data. **A**. No significant effects for any groups on the distance traveled in the center zone relative to the total distance traveled, although a subthreshold increase was observed in the POL-THC group relative to SAL-SAL (t=2.344, p=0.041, q=0.164). No statistically significant differences were observed in the social preference (**B**) or social novelty (**C**) tasks for any of the groups. Finally, no overall differences in prepulse inhibition, based on the maximum startle amplitude were observed (**D**), although a subthreshold interaction was observed between POL-THC vs SAL-SAL and increasing prepulse level (*ns(level, 2)1:group*; t=2.239, p=0.027, q=0.108) (**E**).

### 2.7. Sex differences

#### 2.7.1 Sex differences in response to both prenatal MIA-exposure and adolescent THC exposure

Post-hoc investigation of sex differences in this group revealed a significant three way interaction (*group:ns(age,2)1:sex*) (t=4.238, 5%FDR) (**supplementary figure 2**). In the somatosensory cortex, medial amygdala, ventromedial thalamus, ventral tegmental area (VTA), pontine nucleus, and medulla, male POL-THC offspring had a flatter trajectory, with smaller volumes in the PND 50-85 period relative to SAL-SAL, whereas the opposite pattern was observed in females in all regions except for the medial amygdala and pontine nucleus, where female curves between POL-THC and SAL-SAL offspring were very similar.

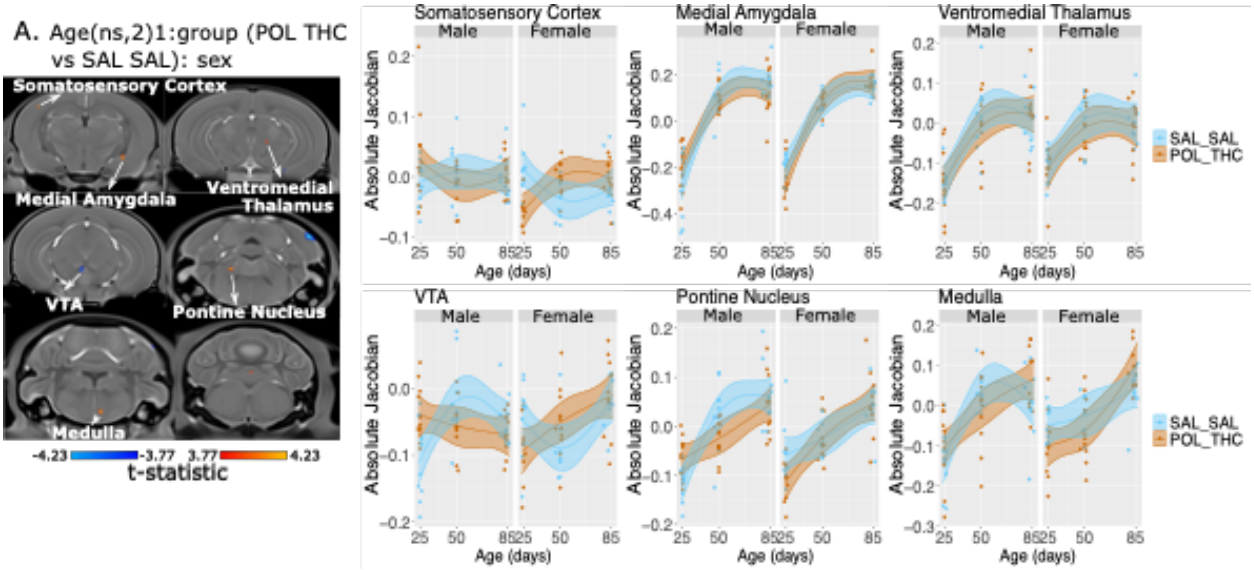

**Supplementary Figure S5.** Sex differences in neuroanatomical alteration due to combined prenatal MIA-exposure and adolescent THC-exposure. **A**. t-statistic map of group (SAL THC vs SAL SAL) by age (first order natural spline of age) by sex thresholded between 5% FDR (top, t=4.23) and 10% FDR (bottom, t=3.77). **B**. Plot of peak voxels selected from regions of interest highlighted in **A**, wherein age is plotted on the x-axis, and the absolute Jacobian determinants plotted on the y-axis. Trajectories reflect the statistical model used (see **2.4.2**).

### 2.6.2 No significant sex differences in behaviour following prenatal MIA-exposure and adolescent THC exposure

Post-hoc investigation of sex differences in this group revealed no significant sex-by-group interactions surviving multiple comparisons correction. A subthreshold sex-by-group interaction (SAL-THC vs. SAL-SAL) was observed in PPI, over increasing prepulse tone (t=2.332, p=0.022, q=0.088), wherein SAL-THC males were impaired in PPI.

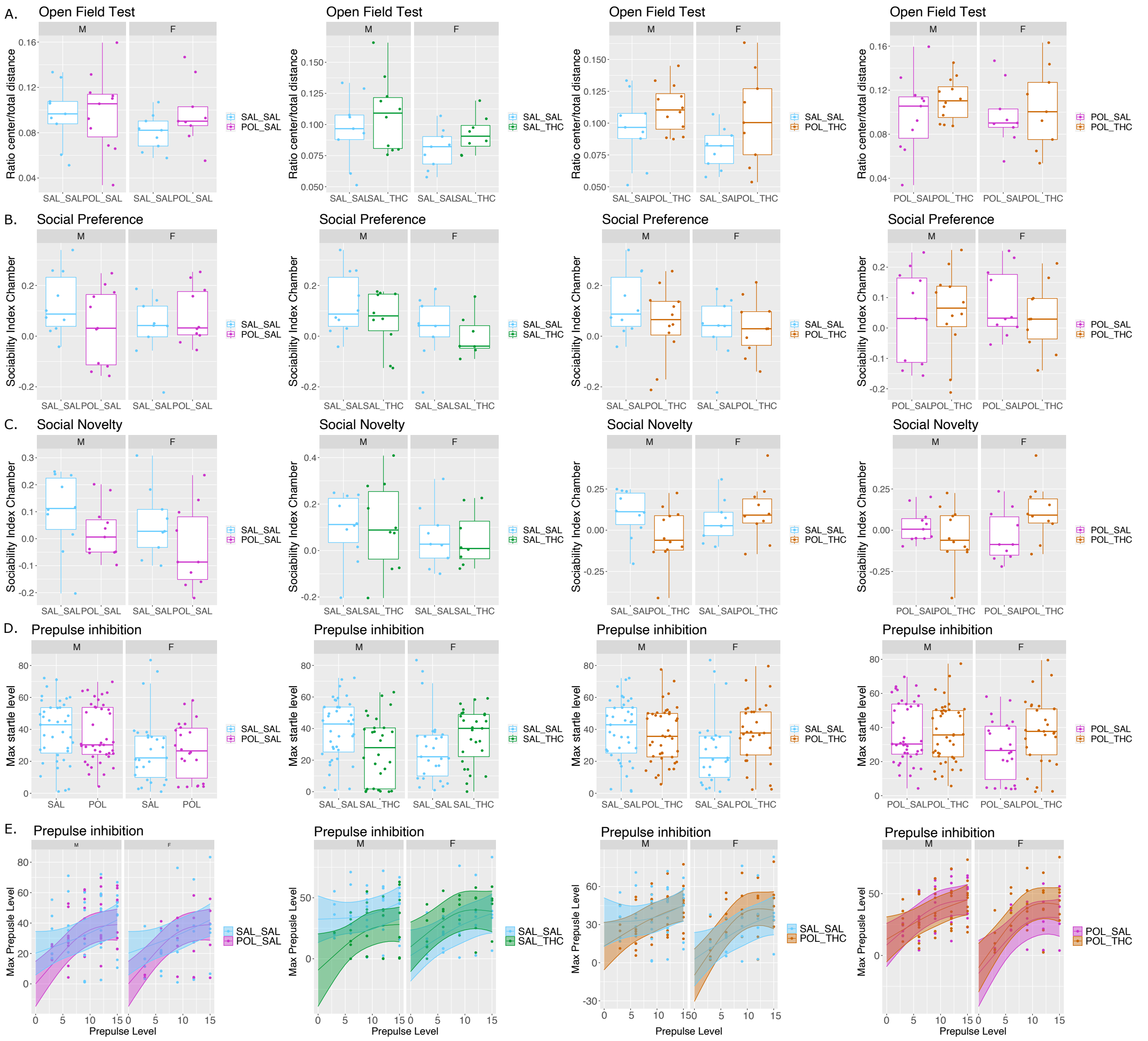

**Supplementary Figure S6.** Sex differences in behaviour due to prenatal MIA-exposure and/or adolescent THC exposure do not affect adult behaviour. Behavioural results for all treatment groups: SAL-SAL (cyan), POL-SAL (magenta), SAL-THC (green), POL-THC (orange). For each boxplot, males (M) are on the left, and females (F) on the right; the midline represents the median of the data, the box represents the interquartile range, with whiskers denoting the full range of the data. No significant effects for any groups on the distance traveled in the center zone relative to the total distance traveled (**A**). No statistically significant differences were observed in the social preference (**B**) or social novelty (**C**) tasks for any of the groups, although a subthreshold sex-by-group (POL-THC vs POL-SAL) interaction was observed in social novelty ($t= 2.194$, $p=0.034$, $q=0.136$) wherein POL-SAL females showed less preference for the novel mouse. No overall differences in prepulse inhibition, based on the maximum startle amplitude were observed (**D**) apart from a subthreshold sex-by-group (POL-THC vs POL-SAL) interaction ($t= 2.194$, $p=0.034$, $q=0.136$) wherein POL-SAL females showed impairment in PPI. Similarly, no group differences were observed over increasing prepulse told, apart from a subthreshold group-by-sex-by-prepulse tone interaction (POL-THC vs SAL-SAL: *sex:ns(level, 2)1:group:sex*; t=2.332, p=0.022, q=0.088) wherein females showed improvement in PPI for louder prepulse tones (**E**).

**Supplementary Table 4. Summary of all behavioural results for all group by sex interactions.** T-values, p-values (uncorrected) and q-values (corrected) are bolded if they survive Bonferroni correction (q-value=p<0.0125).

| | POL-SAL vs. SAL-SAL:sex | SAL-THC vs. SAL-SAL:sex | POL-THC vs. SAL-SAL:sex | POL-SAL vs. POL-THC:sex |
|---|---|---|---|---|
| **OFT** | t=0.919, p=0.366, q=1.000 | t=-0.005, p=0.996, q=1.000 | t=0.529, p=0.601, q=1.000 | t=-0.400, p=0.692, q=1.000 |
| **SOPT** | t=1.678, p=0.105, q=0.42 | t=0.370, p=0.714, q=1.000 | t=0.952, p=0.347, q=1.000 | t=-0.781, p=0.440, q=1.000 |
| **SONT** | t=-0.201, p=0.842, q=1.000 | t=-0.133, p=0.895, q=1.000 | t=1.886, p=0.067, q=0.268 | t= 2.194, p=0.034, q=0.136 |
| **PPI** | **Overall:** t=0.169, p=0.867, q=1.000 **By PP tone (ns(level, 2)1:group:sex)** t=-0.560, p=0.577, q=1.000 | **Overall:** t= 1.841, p=0.079, q=0.316 **By PP tone (ns(level, 2)1:group:sex)** t=-1.331, p=0.187, q=0.748 | **Overall:** t=1.458, p=0.162, q=0.648 **By PP tone (ns(level, 2)1:group:sex)** t=2.332, p=0.022, q=0.088 | **Overall:** t=1.751, p=0.092, q=0.368 **By PP tone (ns(level, 2)1:group:sex)** t=0.723, p=0.471, q=1.000 |

### 2.7 PLS of within subject volume change - behaviour

PLS was used to examine within subject volume change from the post-treatment timepoint (PND50) to the adult timepoint (PND85) at a voxel level across the brain and the same 18 behavioural and demographics metrics as in **3.5.1**. We identified two significant LV (LV1: p<0.00001, %covariance=24%; LV2: p=0.01, %covariance=19%; **supplementary figure S4)**.

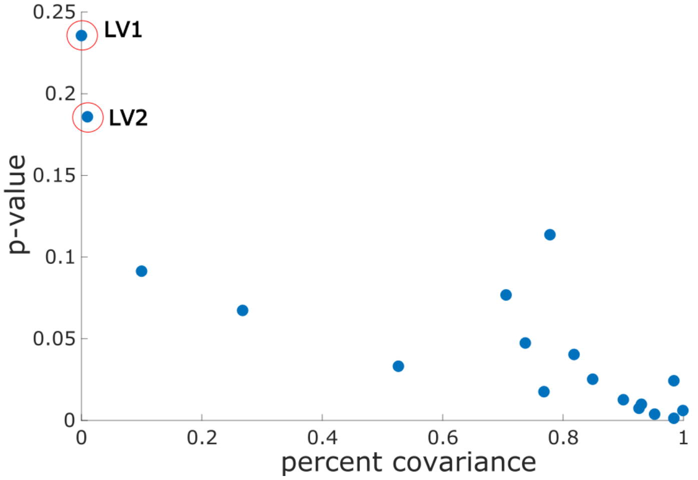

**Supplementary Figure S7.** Covariance explained (y-axis) and permutation p-values (x-axis) for all 18 LVs in the PLS analysis based on the difference in volume from PND 50 to 85, and behaviour. LV1 and LV2 are circled in red (LV1: $p<0.00001$, %covariance=24%; LV2: $p=0.01$, %covariance=19%).

### 2.8 Risk factor exposure significantly decreases CB1-IR and CB2-IR expression in the brain

In the ACC a significant pre- by postnatal treatment interaction was observed in for optical density measures for both CB1-IR intensity ($F(5,40)=14.408$, $p=0.000489$) and CB2-IR intensity ($F(5,40)=4.612$, $p= 4.04e-05$). The same pattern was observed in the SMCTX (CB1: $F(5,40)=2.221$, $p=0.03208$; CB2: $F(5,40)=4.495$, $p=5.82e-05$) and the DG molecular layer of the hippocampus (CB1: $F(5,40)=7.522$, $p=0.0091$; CB2: $F(5,40)=3.747$, $p= 0.000552$). In the ACC, Tukey's post hoc test revealed a significant decrease in CB1-IR optical intensity for SAL-THC ($p<0.05$) and POL-SAL ($p<0.00001$) relative to SAL-SAL, while CB2-IR optical intensity was reduced across all three risk factor groups ($p<0.014$). In the SMCTX, the POL-SAL groups had reduced CB1-IR optical intensity relative to SAL-SAL ($p=0.034$), while all 3 risk factor groups had reduced CB2 intensity relative to SAL-SAL ($p<0.005$). In the DG molecular layer optical density was reduced in all three risk factor groups for CB1-IR ($p<0.024$) and CB2-IR ($p<0.001$) relative to SAL-SAL.

**Supplementary Table 5.** CB1-IR cell (neuron) density in the ACC ANCOVA results & Tukey's post-hoc test (model: Prenatal treatment (TX) * Adolescent treatment (TX) + Sex + Hemisphere)

| ANCOVA | Df | Sum Sq | Mean Sq | F value | Pr(>F) |
|---|---|---|---|---|---|
| Prenatal_TX | 1 | 1.402 | 1.402 | 11.338 | 0.00169 |
| Adolescent_TX | 1 | 2.382 | 2.382 | 19.264 | 0.00008 |
| Sex | 1 | 0.028 | 0.028 | 0.224 | 0.63826 |
| Hemisphere | 1 | 0.003 | 0.003 | 0.021 | 0.88665 |
| Prenatal_TX:Adolescent_TX | 1 | 1.564 | 1.564 | 12.651 | 0.00098 |
| Residuals | 40 | 4.947 | 0.1237 | | |
| **TUKEY'S** | **Difference** | **Lower** | **Upper** | **p adj** | |
| Prenatal_TX (POL-SAL) | -0.350 | -0.559 | -0.140 | 0.002 | * |
| Adolescent_TX (THC-SAL) | -0.455 | -0.665 | -0.245 | <0.00001 | *** |
| Sex (F-M) | 0.049 | -0.163 | 0.260 | 0.645 | |
| Hemisphere (R-L) | -0.015 | -0.224 | 0.195 | 0.887 | |
| POL:SAL-SAL:SAL | -0.753 | -1.157 | -0.350 | <0.00001 | *** |
| SAL:THC-SAL:SAL | -0.839 | -1.243 | -0.435 | <0.00001 | *** |
| POL:THC-SAL:SAL | -0.861 | -1.265 | -0.457 | <0.00001 | *** |
| SAL:THC-POL:SAL | -0.086 | -0.471 | 0.299 | 0.932 | |
| POL:THC-POL:SAL | -0.108 | -0.493 | 0.277 | 0.876 | |
| POL:THC-SAL:THC | -0.022 | -0.407 | 0.363 | 0.999 | |

**Supplementary Table 6.** CB1-IR IR fiber density (optical intensity) in the ACC ANCOVA results & Tukey's post-hoc test (model: Prenatal treatment (TX) * Adolescent treatment (TX) + Sex + Hemisphere)

| ANCOVA | Df | Sum Sq | Mean Sq | F value | Pr(>F) |
|---|---|---|---|---|---|
| Prenatal_TX | 1 | 424 | 424 | 7.261 | 0.01025 |
| Adolescent_TX | 1 | 1.4 | 1.4 | 0.024 | 0.87675 |
| Sex | 1 | 440.3 | 440.3 | 7.54 | 0.00900 |
| Hemisphere | 1 | 43.8 | 43.8 | 0.751 | 0.39139 |
| Prenatal_TX:Adolescent_TX | 1 | 841.4 | 841.4 | 14.408 | 0.00049 |
| Residuals | 40 | 4.947 | 0.1237 | | |
| **TUKEY'S** | **Difference** | **Lower** | **Upper** | **p adj** | |
| Prenatal_TX (POL-SAL) | -6.078 | -10.637 | -1.519 | 0.010 | * |
| Adolescent_TX (THC-SAL) | -0.352 | -4.910 | 4.207 | 0.877 | |
| Sex (F-M) | -6.116 | -10.710 | -1.523 | 0.010 | * |

| | | | | | |
|---|---|---|---|---|---|
| Hemisphere (R-L) | -1.953 | -6.507 | 2.602 | 0.391 | |
| POL:SAL-SAL:SAL | -14.976 | -23.747 | -6.206 | <0.00001 | *** |
| SAL:THC-SAL:SAL | -9.234 | -18.005 | -0.464 | 0.036 | * |
| POL:THC-SAL:SAL | -7.254 | -16.025 | 1.517 | 0.136 | |
| SAL:THC-POL:SAL | 5.742 | -2.621 | 14.104 | 0.270 | |
| POL:THC-POL:SAL | 7.722 | -0.640 | 16.085 | 0.079 | |
| POL:THC-SAL:THC | 1.980 | -6.382 | 10.343 | 0.920 | |

**Supplementary Table 7.** CB2-IR cell (neuron) density in the ACC ANCOVA results & Tukey's post-hoc test (model Prenatal treatment (TX) * Adolescent treatment (TX) + Sex + Hemisphere)

| **ANCOVA** | **Df** | **Sum Sq** | **Mean Sq** | **F value** | **Pr(>F)** |
|---|---|---|---|---|---|
| Prenatal_TX | 1 | 1.284 | 1.2836 | 5.031 | 0.0305 |
| Adolescent_TX | 1 | 2.342 | 2.3422 | 9.181 | 0.00427 |
| Sex | 1 | 0.251 | 0.2513 | 0.985 | 0.32688 |
| Hemisphere | 1 | 0.043 | 0.0428 | 0.168 | 0.68427 |
| Prenatal_TX:Adolescent_TX | 1 | 1.544 | 1.5437 | 6.051 | 0.01832 |
| Residuals | 40 | 10.205 | 0.2551 | | |
| **TUKEY'S** | **Difference** | **Lower** | **Upper** | **p adj** | |
| Prenatal_TX (POL-SAL) | -0.334 | -0.636 | -0.033 | 0.031 | * |
| Adolescent_TX (THC-SAL) | -0.451 | -0.753 | -0.150 | 0.004 | * |
| Sex (F-M) | 0.146 | -0.157 | 0.450 | 0.337 | |
| Hemisphere (R-L) | 0.061 | -0.240 | 0.362 | 0.684 | |
| POL:SAL-SAL:SAL | -0.735 | -1.315 | -0.156 | 0.008 | * |
| SAL:THC-SAL:SAL | -0.833 | -1.412 | -0.253 | 0.002 | * |
| POL:THC-SAL:SAL | -0.842 | -1.421 | -0.262 | 0.002 | * |
| SAL:THC-POL:SAL | -0.097 | -0.650 | 0.455 | 0.965 | |
| POL:THC-POL:SAL | -0.106 | -0.659 | 0.446 | 0.955 | |
| POL:THC-SAL:THC | -0.009 | -0.562 | 0.544 | 1.000 | |

**Supplementary Table 8.** CB2-IR IR fiber density (optical intensity) in the ACC ANCOVA results & Tukey's post-hoc test (model: Prenatal treatment (TX) * Adolescent treatment (TX) + Sex + Hemisphere)

| **ANCOVA** | **Df** | **Sum Sq** | **Mean Sq** | **F value** | **Pr(>F)** |
|---|---|---|---|---|---|
| Prenatal_TX | 1 | 482.5 | 482.5 | 9.438 | 0.00381 |
| Adolescent_TX | 1 | 60.7 | 60.7 | 1.188 | 0.28218 |
| Sex | 1 | 101.6 | 101.6 | 1.988 | 0.16632 |

| | | | | | |
|---|---|---|---|---|---|
| Hemisphere | 1 | 27.4 | 27.4 | 0.537 | 0.46803 |
| Prenatal_TX:Adolescent_TX | 1 | 1087.3 | 1087.3 | 21.271 | 4.04E-05 |
| Residuals | 40 | 2044.7 | 51.1 | | |
| **TUKEY'S** | **Difference** | **Lower** | **Upper** | **p adj** | |
| Prenatal_TX (POL-SAL) | -6.483 | -10.748 | -2.218 | 0.004 | * |
| Adolescent_TX (THC-SAL) | -2.298 | -6.563 | 1.967 | 0.283 | |
| Sex (F-M) | -2.938 | -7.236 | 1.360 | 0.175 | |
| Hemisphere (R-L) | -1.545 | -5.806 | 2.716 | 0.468 | |
| POL:SAL-SAL:SAL | -16.685 | -24.890 | -8.479 | <0.00001 | *** |
| SAL:THC-SAL:SAL | -12.400 | -20.605 | -4.194 | 0.001 | ** |
| POL:THC-SAL:SAL | -9.809 | -18.014 | -1.603 | 0.014 | * |
| SAL:THC-POL:SAL | 4.285 | -3.539 | 12.109 | 0.466 | |
| POL:THC-POL:SAL | 6.876 | -0.948 | 14.700 | 0.103 | |
| POL:THC-SAL:THC | 2.591 | -5.233 | 10.415 | 0.811 | |

**Supplementary Table 9.** CB1-IR cell (neuron) density in the SMCTX ANCOVA results & Tukey's post-hoc test (model: Prenatal treatment (TX) * Adolescent treatment (TX) + Sex + Hemisphere)

| **ANCOVA** | **Df** | **Sum Sq** | **Mean Sq** | **F value** | **Pr(>F)** |
|---|---|---|---|---|---|
| Prenatal_TX | 1 | 1.0958 | 1.0958 | 18.681 | 9.95E-05 |
| Adolescent_TX | 1 | 1.3719 | 1.3719 | 23.389 | 1.99E-05 |
| Sex | 1 | 0.0331 | 0.0331 | 0.564 | 0.457 |
| Hemisphere | 1 | 0.051 | 0.051 | 0.87 | 0.357 |
| Prenatal_TX:Adolescent_TX | 1 | 2.4364 | 2.4364 | 41.537 | 1.12E-07 |
| Residuals | 40 | 2.3463 | 0.0587 | | |
| **TUKEY'S** | **Difference** | **Lower** | **Upper** | **p adj** | |
| Prenatal_TX (POL-SAL) | -0.309 | -0.453 | -0.164 | <0.00001 | *** |
| Adolescent_TX (THC-SAL) | -0.345 | -0.490 | -0.201 | <0.00001 | *** |
| Sex (F-M) | 0.053 | -0.093 | 0.199 | 0.466 | |
| Hemisphere (R-L) | -0.067 | -0.211 | 0.078 | 0.357 | |
| POL:SAL-SAL:SAL | -0.803 | -1.081 | -0.525 | <0.00001 | *** |
| SAL:THC-SAL:SAL | -0.824 | -1.102 | -0.546 | <0.00001 | *** |
| POL:THC-SAL:SAL | -0.714 | -0.992 | -0.436 | <0.00001 | *** |
| SAL:THC-POL:SAL | -0.021 | -0.286 | 0.244 | 0.996 | |
| POL:THC-POL:SAL | 0.088 | -0.177 | 0.353 | 0.808 | |

| | | | | |
|---|---|---|---|---|
| POL:THC-SAL:THC | 0.110 | -0.155 | 0.375 | 0.685 |

**Supplementary Table 10.** CB1-IR IR fiber density (optical intensity) in the SMCTX ANCOVA results & Tukey's post-hoc test (model: Prenatal treatment (TX) * Adolescent treatment (TX) + Sex + Hemisphere)

| **ANCOVA** | **Df** | **Sum Sq** | **Mean Sq** | **F value** | **Pr(>F)** |
|---|---|---|---|---|---|
| Prenatal_TX | 1 | 434 | 433.7 | 3.30E+00 | 7.66E-02 |
| Adolescent_TX | 1 | 6 | 6.3 | 0.048 | 0.8277 |
| Sex | 1 | 225 | 225 | 1.713 | 0.198 |
| Hemisphere | 1 | 0 | 0.4 | 0.003 | 0.9559 |
| Prenatal_TX:Adolescent_TX | 1 | 648 | 647.6 | 4.933 | 0.0321 |
| Residuals | 40 | 5252 | 131.3 | | |
| **TUKEY'S** | **Difference** | **Lower** | **Upper** | **p adj** | |
| Prenatal_TX (POL-SAL) | -6.147 | -12.982 | 0.688 | 0.077 | |
| Adolescent_TX (THC-SAL) | -0.740 | -7.575 | 6.100 | 0.828 | |
| Sex (F-M) | -4.372 | -11.260 | 2.516 | 0.207 | |
| Hemisphere (R-L) | -0.188 | -7.017 | 6.641 | 0.956 | |
| POL:SAL-SAL:SAL | -13.973 | -27.124 | -0.823 | 0.034 | * |
| SAL:THC-SAL:SAL | -8.534 | -21.684 | 4.616 | 0.317 | |
| POL:THC-SAL:SAL | -7.631 | -20.781 | 5.520 | 0.415 | |
| SAL:THC-POL:SAL | 5.439 | -7.099 | 17.978 | 0.653 | |
| POL:THC-POL:SAL | 6.342 | -6.196 | 18.881 | 0.534 | |
| POL:THC-SAL:THC | 0.903 | -11.635 | 13.442 | 0.997 | |

**Supplementary Table 11.** CB2-IR cell (neuron) density in the SMCTX ANCOVA results & Tukey's post-hoc test (model: Prenatal treatment (TX) * Adolescent treatment (TX) + Sex + Hemisphere)

| **ANCOVA** | **Df** | **Sum Sq** | **Mean Sq** | **F value** | **Pr(>F)** |
|---|---|---|---|---|---|
| Prenatal_TX | 1 | 0.837 | 0.837 | 13.243 | 0.001 |
| Adolescent_TX | 1 | 1.113 | 1.113 | 17.601 | <0.00001 |
| Sex | 1 | 0.068 | 0.068 | 1.076 | 0.306 |
| Hemisphere | 1 | 0.003 | 0.003 | 0.045 | 0.834 |
| Prenatal_TX:Adolescent_TX | 1 | 0.937 | 0.937 | 14.823 | <0.00001 |
| Residuals | 40 | 2.5288 | 0.0632 | | |

| TUKEY'S | Difference | Lower | Upper | p adj | |
|---|---|---|---|---|---|
| Prenatal_TX (POL-SAL) | -0.270 | -0.420 | -0.120 | 0.001 | ** |
| Adolescent_TX (THC-SAL) | -0.311 | -0.461 | -0.161 | <0.00001 | *** |
| Sex (F-M) | 0.076 | -0.075 | 0.227 | 0.315 | |
| Hemisphere (R-L) | -0.016 | -0.166 | 0.134 | 0.834 | |
| POL:SAL-SAL:SAL | -0.581 | -0.869 | -0.292 | <0.00001 | *** |
| SAL:THC-SAL:SAL | -0.608 | -0.897 | -0.320 | <0.00001 | *** |
| POL:THC-SAL:SAL | -0.623 | -0.911 | -0.334 | <0.00001 | *** |
| SAL:THC-POL:SAL | -0.027 | -0.303 | 0.248 | 0.993 | |
| POL:THC-POL:SAL | -0.042 | -0.317 | 0.233 | 0.976 | |
| POL:THC-SAL:THC | -0.015 | -0.290 | 0.260 | 0.999 | |

**Supplementary Table 12.** CB2-IR IR fiber density (optical intensity) in the SMCTX ANCOVA results & Tukey's post-hoc test (model: Prenatal treatment (TX) * Adolescent treatment (TX) + Sex + Hemisphere)

| ANCOVA | Df | Sum Sq | Mean Sq | F value | Pr(>F) |
|---|---|---|---|---|---|
| Prenatal_TX | 1 | 580 | 5.80E+02 | 5.933 | 1.94E-02 |
| Adolescent_TX | 1 | 489 | 489.5 | 5.009 | 3.09E-02 |
| Sex | 1 | 328 | 328 | 3.357 | 0.0744 |
| Hemisphere | 1 | 0 | 0 | 0 | 0.9882 |
| Prenatal_TX:Adolescent_TX | 1 | 1975 | 1974.5 | 2.02E+01 | 5.82E-05 |
| Residuals | 40 | 3909 | 97.7 | | |
| **TUKEY'S** | **Difference** | **Lower** | **Upper** | **p adj** | |
| Prenatal_TX (POL-SAL) | -7.107 | -13.004 | -1.210 | 0.019 | * |
| Adolescent_TX (THC-SAL) | -6.523 | -12.420 | -0.626 | 0.031 | * |
| Sex (F-M) | -5.279 | -11.221 | 0.663 | 0.080 | |
| Hemisphere (R-L) | 0.043 | -5.848 | 5.935 | 0.988 | |
| POL:SAL-SAL:SAL | -21.010 | -32.356 | -9.665 | <0.00001 | *** |
| SAL:THC-SAL:SAL | -20.143 | -31.488 | -8.798 | <0.00001 | *** |
| POL:THC-SAL:SAL | -15.178 | -26.523 | -3.833 | 0.005 | ** |
| SAL:THC-POL:SAL | 0.867 | -9.950 | 11.685 | 0.996 | |
| POL:THC-POL:SAL | 5.832 | -4.985 | 16.650 | 0.479 | |
| POL:THC-SAL:THC | 4.965 | -5.853 | 15.782 | 0.612 | |

**Supplementary Table 13.** CB1-IR cell (neuron) density in the DG molecular layer ANCOVA results & Tukey's post-hoc test (model: Prenatal treatment (TX) * Adolescent treatment (TX) + Sex + Hemisphere)

| ANCOVA | Df | Sum Sq | Mean Sq | F value | Pr(>F) |
|---|---|---|---|---|---|
| Prenatal_TX | 1 | 4.11 | 4.11 | 6.235 | 0.0167 |
| Adolescent_TX | 1 | 0.232 | 0.232 | 0.352 | 0.5565 |
| Sex | 1 | 0.29 | 0.29 | 0.44 | 0.5110 |
| Hemisphere | 1 | 0.002 | 0.002 | 0.003 | 0.9590 |
| Prenatal_TX:Adolescent_TX | 1 | 2.654 | 2.654 | 4.027 | 0.0516 |
| Residuals | 40 | 26.366 | 0.659 | | |
| **TUKEY'S** | **Difference** | **Lower** | **Upper** | **p adj** | |
| Prenatal_TX (POL-SAL) | -0.5984 | -1.0827 | -0.1140 | 0.0167 | |
| Adolescent_TX (THC-SAL) | 0.1419 | -0.3424 | 0.6263 | 0.5570 | |
| Sex (F-M) | 0.1571 | -0.3310 | 0.6451 | 0.5191 | |
| Hemisphere (R-L) | 0.0124 | -0.4715 | 0.4962 | 0.9591 | |
| POL:SAL-SAL:SAL | -1.0907 | -2.0225 | -0.1590 | 0.0162 | * |
| SAL:THC-SAL:SAL | -0.3566 | -1.2884 | 0.5752 | 0.7354 | |
| POL:THC-SAL:SAL | -0.4950 | -1.4268 | 0.4368 | 0.4922 | |
| SAL:THC-POL:SAL | 0.7341 | -0.1543 | 1.6226 | 0.1366 | |
| POL:THC-POL:SAL | 0.5957 | -0.2927 | 1.4842 | 0.2896 | |
| POL:THC-SAL:THC | -0.1384 | -1.0268 | 0.7500 | 0.9752 | |

**Supplementary Table 14.** CB1-IR IR fiber density (optical intensity) in the DG molecular layer ANCOVA results & Tukey's post-hoc test (model: Prenatal treatment (TX) * Adolescent treatment (TX) + Sex + Hemisphere)

| ANCOVA | Df | Sum Sq | Mean Sq | F value | Pr(>F) |
|---|---|---|---|---|---|
| Prenatal_TX | 1 | 107.8 | 107.79 | 9.66E+00 | 0.00345 |
| Adolescent_TX | 1 | 24.6 | 24.56 | 2.202 | 0.14566 |
| Sex | 1 | 1.8 | 1.77 | 0.158 | 0.69266 |
| Hemisphere | 1 | 0.2 | 0.24 | 0.022 | 0.88400 |
| Prenatal_TX:Adolescent_TX | 1 | 83.9 | 83.9 | 7.522 | 0.00907 |
| Residuals | 40 | 446.1 | 11.15 | | |
| **TUKEY'S** | **Difference** | **Lower** | **Upper** | **p adj** | |
| Prenatal_TX (POL-SAL) | -3.064 | -5.057 | -1.072 | 0.003 | * |
| Adolescent_TX (THC-SAL) | -1.461 | -3.454 | 0.531 | 0.146 | |

| | | | | | |
|---|---|---|---|---|---|
| Sex (F-M) | 0.388 | -1.620 | 2.395 | 0.698 | |
| Hemisphere (R-L) | 0.145 | -1.845 | 2.135 | 0.884 | |
| POL:SAL-SAL:SAL | -5.936 | -9.769 | -2.103 | 0.001 | ** |
| SAL:THC-SAL:SAL | -4.269 | -8.102 | -0.436 | 0.024 | * |
| POL:THC-SAL:SAL | -4.850 | -8.683 | -1.017 | 0.008 | * |
| SAL:THC-POL:SAL | 1.667 | -1.988 | 5.321 | 0.617 | |
| POL:THC-POL:SAL | 1.085 | -2.569 | 4.740 | 0.856 | |
| POL:THC-SAL:THC | -0.581 | -4.236 | 3.073 | 0.974 | |

**Supplementary Table 15.** CB2-IR cell (neuron) density in the DG molecular layer ANCOVA results & Tukey's post-hoc test (model: Prenatal treatment (TX) * Adolescent treatment (TX) + Sex + Hemisphere)

| **ANCOVA** | **Df** | **Sum Sq** | **Mean Sq** | **F value** | **Pr(>F)** |
|---|---|---|---|---|---|
| Prenatal_TX | 1 | 15.75 | 15.75 | 11.063 | 0.00190 |
| Adolescent_TX | 1 | 7.15 | 7.153 | 5E+00 | 0.03061 |
| Sex | 1 | 0.54 | 0.543 | 0.382 | 0.54028 |
| Hemisphere | 1 | 0.1 | 0.097 | 0.068 | 0.79513 |
| Prenatal_TX:Adolescent_TX | 1 | 25.35 | 25.352 | 17.807 | 0.00014 |
| Residuals | 40 | 56.95 | 1.424 | | |
| **TUKEY'S** | **Difference** | **Lower** | **Upper** | **p adj** | |
| Prenatal_TX (POL-SAL) | -1.171 | -1.883 | -0.460 | 0.002 | * |
| Adolescent_TX (THC-SAL) | -0.789 | -1.500 | -0.077 | 0.031 | * |
| Sex (F-M) | 0.215 | -0.502 | 0.932 | 0.548 | |
| Hemisphere (R-L) | -0.092 | -0.803 | 0.619 | 0.795 | |
| POL:SAL-SAL:SAL | -2.749 | -4.118 | -1.380 | <0.00001 | *** |
| SAL:THC-SAL:SAL | -2.332 | -3.701 | -0.963 | <0.00001 | *** |
| POL:THC-SAL:SAL | -2.138 | -3.507 | -0.768 | 0.001 | ** |
| SAL:THC-POL:SAL | 0.417 | -0.889 | 1.723 | 0.827 | |
| POL:THC-POL:SAL | 0.611 | -0.694 | 1.917 | 0.596 | |
| POL:THC-SAL:THC | 0.194 | -1.111 | 1.500 | 0.978 | |

**Supplementary Table 16.** CB2-IR IR fiber density (optical intensity)in the DG molecular layer ANCOVA results & Tukey's post-hoc test (Prenatal treatment (TX) * Adolescent treatment (TX) + Sex + Hemisphere)

| ANCOVA | Df | Sum Sq | Mean Sq | F value | Pr(>F) |
|---|---|---|---|---|---|
| Prenatal_TX | 1 | 183.3 | 183.3 | 11.153 | 0.0018 |
| Adolescent_TX | 1 | 78.3 | 78.33 | 4.766 | 0.0350 |
| Sex | 1 | 4.3 | 4.27 | 2.60E-01 | 0.6130 |
| Hemisphere | 1 | 0 | 0.04 | 0.002 | 0.9616 |
| Prenatal_TX:Adolescent_TX | 1 | 225.2 | 225.16 | 13.699 | 0.0006 |
| Residuals | 40 | 657.4 | 16.44 | | |
| **TUKEY'S** | **Difference** | **Lower** | **Upper** | **p adj** | |
| Prenatal_TX (POL-SAL) | -3.996 | -6.415 | -1.578 | 0.002 | ** |
| Adolescent_TX (THC-SAL) | -2.610 | -5.028 | -0.191 | 0.035 | * |
| Sex (F-M) | 0.602 | -1.835 | 3.039 | 0.620 | |
| Hemisphere (R-L) | -0.058 | -2.474 | 2.358 | 0.962 | |
| POL:SAL-SAL:SAL | -8.710 | -13.363 | -4.057 | <0.00001 | *** |
| SAL:THC-SAL:SAL | -7.210 | -11.863 | -2.557 | 0.001 | ** |
| POL:THC-SAL:SAL | -7.148 | -11.801 | -2.495 | 0.001 | ** |
| SAL:THC-POL:SAL | 1.500 | -2.936 | 5.936 | 0.802 | |
| POL:THC-POL:SAL | 1.562 | -2.875 | 5.998 | 0.782 | |
| POL:THC-SAL:THC | 0.062 | -4.375 | 4.498 | 1.000 | |

**Supplementary Table 17.** CB1- IR fiber density (optical intensity)in the STR ANCOVA results & Tukey's post-hoc test (model: Prenatal treatment (TX) * Adolescent treatment (TX) + Sex + Hemisphere)

| ANCOVA | Df | Sum Sq | Mean Sq | F value | Pr(>F) |
|---|---|---|---|---|---|
| Prenatal_TX | 1 | 600 | 600 | 8.54 | 0.0057 |
| Adolescent_TX | 1 | 105.4 | 105.4 | 1.50E+00 | 0.2277 |
| Sex | 1 | 2.9 | 2.9 | 0.042 | 0.8391 |
| Hemisphere | 1 | 4 | 4 | 0.057 | 0.8122 |
| Prenatal_TX:Adolescent_TX | 1 | 1507.1 | 1507.1 | 21.452 | <0.00001 |
| Residuals | 40 | 2810.2 | 70.3 | | |
| **TUKEY'S** | **Difference** | **Lower** | **Upper** | **p adj** | |
| Prenatal_TX (POL-SAL) | -7.230 | -12.230 | -2.230 | 0.006 | ** |
| Adolescent_TX (THC-SAL) | -3.028 | -8.028 | 1.972 | 0.228 | |
| Sex (F-M) | 0.499 | -4.539 | 5.538 | 0.842 | |
| Hemisphere (R-L) | -0.591 | -5.587 | 4.404 | 0.812 | |

| | | | | | |
|---|---|---|---|---|---|
| POL:SAL-SAL:SAL | -19.255 | -28.875 | -9.635 | <0.00001 | *** |
| SAL:THC-SAL:SAL | -14.921 | -24.541 | -5.301 | 0.001 | ** |
| POL:THC-SAL:SAL | -11.482 | -21.102 | -1.863 | 0.014 | * |
| SAL:THC-POL:SAL | 4.334 | -4.838 | 13.506 | 0.589 | |
| POL:THC-POL:SAL | 7.773 | -1.399 | 16.945 | 0.122 | |
| POL:THC-SAL:THC | 3.439 | -5.733 | 12.611 | 0.747 | |

**Supplementary Table 18.** CB2-IR IR fiber density (optical intensity)in the STR ANCOVA results & Tukey's post-hoc test (model: Prenatal treatment (TX) * Adolescent treatment (TX) + Sex + Hemisphere)

| **ANCOVA** | **Df** | **Sum Sq** | **Mean Sq** | **F value** | **Pr(>F)** |
|---|---|---|---|---|---|
| Prenatal_TX | 1 | 1727 | 1726.8 | 14.412 | 0.00049 |
| Adolescent_TX | 1 | 241 | 241.1 | 2.012 | 0.16400 |
| Sex | 1 | 1 | 1.1 | 0.009 | 0.92400 |
| Hemisphere | 1 | 5 | 4.8 | 0.04 | 0.84273 |
| Prenatal_TX:Adolescent_TX | 1 | 2403 | 2402.8 | 20.054 | 0.00006 |
| Residuals | 40 | 4793 | 119.8 | | |
| **TUKEY'S** | **Difference** | **Lower** | **Upper** | **p adj** | |
| Prenatal_TX (POL-SAL) | -12.265 | -18.795 | -5.735 | <0.00001 | *** |
| Adolescent_TX (THC-SAL) | -4.579 | -11.109 | 1.951 | 0.164 | |
| Sex (F-M) | -0.307 | -6.887 | 6.273 | 0.925 | |
| Hemisphere (R-L) | -0.645 | -7.168 | 5.879 | 0.843 | |
| POL:SAL-SAL:SAL | -27.483 | -40.046 | -14.921 | <0.00001 | *** |
| SAL:THC-SAL:SAL | -19.598 | -32.160 | -7.035 | 0.001 | ** |
| POL:THC-SAL:SAL | -18.427 | -30.989 | -5.864 | 0.002 | ** |
| SAL:THC-POL:SAL | 7.886 | -4.092 | 19.864 | 0.305 | |
| POL:THC-POL:SAL | 9.057 | -2.921 | 21.035 | 0.195 | |
| POL:THC-SAL:THC | 1.171 | -10.807 | 13.149 | 0.994 | |

**Supplementary Table 19.** CB1-IR IR fiber density (optical intensity)in the CA1 molecular layer ANCOVA results & Tukey's post-hoc test (model: Prenatal_TX * Adolescent_TX + Sex + Hemisphere)

| **ANCOVA** | **Df** | **Sum Sq** | **Mean Sq** | **F value** | **Pr(>F)** |
|---|---|---|---|---|---|
| Prenatal_TX | 1 | 397 | 396.9 | 2.92E+00 | 0.095 |
| Adolescent_TX | 1 | 42 | 42.3 | 0.311 | 0.580 |
| Sex | 1 | 360 | 360.1 | 2.648 | 0.112 |
| Hemisphere | 1 | 29 | 28.8 | 0.212 | 0.648 |
| Prenatal_TX:Adolescent_TX | 1 | 841 | 840.9 | 6.184 | 0.017 |

| Residuals | 40 | 5440 | 136 | | |
|---|---|---|---|---|---|
| **TUKEY'S** | **Difference** | **Lower** | **Upper** | **p adj** | |
| Prenatal_TX (POL-SAL) | -5.881 | -12.837 | 1.076 | 0.095 | |
| Adolescent_TX (THC-SAL) | -1.918 | -8.875 | 5.038 | 0.580 | |
| Sex (F-M) | -5.531 | -12.541 | 1.479 | 0.119 | |
| Hemisphere (R-L) | -1.584 | -8.534 | 5.366 | 0.648 | |
| POL:SAL-SAL:SAL | -14.847 | -28.231 | -1.463 | 0.025 | * |
| SAL:THC-SAL:SAL | -10.801 | -24.185 | 2.582 | 0.151 | |
| POL:THC-SAL:SAL | -8.697 | -22.081 | 4.687 | 0.316 | |
| SAL:THC-POL:SAL | 4.046 | -8.715 | 16.807 | 0.830 | |
| POL:THC-POL:SAL | 6.150 | -6.611 | 18.911 | 0.573 | |
| POL:THC-SAL:THC | 2.104 | -10.657 | 14.865 | 0.971 | |

**Supplementary Table 20.** CB2-IR fiber density (optical intensity) in the CA1 molecular layer ANCOVA results & Tukey's post-hoc test (model: Prenatal treatment (TX) * Adolescent treatment (TX) + Sex + Hemisphere)

| **ANCOVA** | **Df** | **Sum Sq** | **Mean Sq** | **F value** | **Pr(>F)** |
|---|---|---|---|---|---|
| Prenatal_TX | 1 | 744 | 744 | 4.727 | 0.0357 |
| Adolescent_TX | 1 | 2065 | 2065 | 1.31E+01 | 0.0008 |
| Sex | 1 | 80 | 80 | 0.51 | 0.4793 |
| Hemisphere | 1 | 12 | 12 | 0.077 | 0.7823 |
| Prenatal_TX:Adolescent_TX | 1 | 6030 | 6030 | 38.324 | <0.00001 |
| Residuals | 40 | 6294 | 157 | | |
| **TUKEY'S** | **Difference** | **Lower** | **Upper** | **p adj** | |
| Prenatal_TX (POL-SAL) | -8.050 | -15.533 | -0.567 | 0.036 | * |
| Adolescent_TX (THC-SAL) | -13.398 | -20.881 | -5.915 | 0.001 | ** |
| Sex (F-M) | 2.611 | -4.929 | 10.151 | 0.488 | |
| Hemisphere (R-L) | -1.029 | -8.505 | 6.447 | 0.782 | |
| POL:SAL-SAL:SAL | -32.438 | -46.834 | -18.041 | <0.00001 | *** |
| SAL:THC-SAL:SAL | -37.203 | -51.599 | -22.807 | <0.00001 | *** |
| POL:THC-SAL:SAL | -24.247 | -38.644 | -9.851 | <0.00001 | *** |
| SAL:THC-POL:SAL | -4.765 | -18.492 | 8.961 | 0.789 | |
| POL:THC-POL:SAL | 8.190 | -5.536 | 21.917 | 0.391 | |
| POL:THC-SAL:THC | 12.956 | -0.771 | 26.682 | 0.070 | |

### Maternal Immune Activation Model Reporting Guidelines Checklist

| ARRIVE Reporting Guideline & Recommendation | Arrive Item | MIA Model Specific Reporting Recommendation<br>*Please complete this chart for each point outlined below. If not applicable, write N/A* |
|---|---|---|
| **Study design**<br>➢Overview of immune activation issues<br><br>For each experiment, give brief details of the study design including:<br>a. The number of experimental and control groups.<br>b. Any steps taken to minimize the effects of subjective bias when allocating animals to treatment (e.g. randomization procedure) and when assessing results (e.g. if done, describe who was blinded and when).<br>c. The experimental unit (e.g. a single animal, group or cage of animals).<br><br>A time-line diagram or flow chart can be useful to illustrate how complex study designs were carried out. | 6 | MIA Specific Reporting:<br>a. General need for improved reporting in MIA model methods + reporting pilot data<br>○ Details on pilot data: |
| **Experimental procedures**<br>➢Compounds<br>➢Validation measures<br><br>For each experiment and each experimental group, including controls, provide precise details of all procedures carried out. For example:<br>a. How (e.g. drug formulation and dose, site and route of administration, anaesthesia and analgesia used [including monitoring], surgical procedure, method of euthanasia). Provide details of any specialist equipment used, including supplier(s).<br>b. When (e.g. time of day).<br>c. Where (e.g. home cage, laboratory, water maze).<br>d. Why (e.g. rationale for choice of specific anaesthetic, route of administration, drug dose used). | 7 | Provide details of:<br>a. Compounds – source, vehicle, preparation/storage, administration route, volume administered, whether anesthetics were used at time of immune challenge.<br>○ Name of compound:<br>○ Catalogue number:<br>○ Lot number:<br>○ Vehicle control used:<br>○ Route of administration:<br>○ Volume administered:<br>○ Storage conditions:<br>○ Anesthetic (type, dose, duration) used:<br><br>b. Housing variables at injection - temperature of room at injection time, cage change at time of injection or not<br>○ Light cycle of animal housing room:<br>○ Time of day of injection:<br>○ Room temperature at injection time:<br>○ Did a cage change occur at time of injection: |

| | | |
|---|---|---|
| | | c. Validation of immune activation – behavior, physiological indices and/or cytokine data, including pilot dosing data<br>○ Method used to verify immune activation:<br><br>d. Validation of gestational timing – vaginal plug, estrous cycle, weight gain<br>○ Method of validating gestational timing:<br><br>Additional comments: |
| **Experimental animals**<br>➢Species/strain/vendor<br><br>a. Provide details of the animals used, including species, strain, sex, developmental stage (e.g. mean or median age plus age range) and weight (e.g. mean or median weight plus weight range).<br>b. Provide further relevant information such as the source of animals, international strain nomenclature, genetic modification status (e.g. knock-out or transgenic), genotype, health/immune status, drug or test naïve, previous procedures, etc. | 8 | Provide details of:<br>a. Species – considerations for appropriate species (mouse, rat, non human primate, other)<br>○ Species:<br><br>b. Strain – variability in strain can influence model<br>○ Strain:<br><br>c. Maternal/Offspring Physiological Variables at time of immune challenge – age, body weight<br>○ Maternal Age at challenge:<br>○ Maternal Body weight:<br>○ Offspring Age at challenge:<br>○ Offspring Sex:<br>○ Offspring Body weight:<br><br>d. Vendor – even within the same strain, vendor can influence endpoints<br>○ Vendor:<br>○ Location of Vendor:<br>○ Room/area where animals originated from: |

| | | |
|---|---|---|
| | | Additional Comments: |
| **Housing and husbandry**<br>➢Cage, ventilation, bedding, enrichment<br><br>Provide details of:<br>a. Housing (type of facility e.g. specific pathogen free [SPF]; type of cage or housing; bedding material; number of cage companions; tank shape and material etc. for fish).<br>b. Husbandry conditions (e.g. breeding program, light/dark cycle, temperature, quality of water etc for fish, type of food, access to food and water, environmental enrichment).<br>c. Welfare-related assessments and interventions that were carried out prior to, during, or after the experiment. | 9 | Provide details of:<br>a. Caging systems<br>○ *At breeding*<br>Material of cage:<br>Cage dimensions:<br>○ *After parturition*<br>Material of cage:<br>Cage dimensions:<br>○ *At weaning*<br>Material of cage:<br>Cage dimensions:<br><br>b. Animal Holding room<br>○ Temperature in room:<br>○ Humidity in room:<br>○ Ventilation system:<br>○ Specific pathogen free [SPF]:<br>○ Are males & females housed in the same or separate rooms:<br><br>c. Bedding exchanges/bedding type<br>○ Type of cage bedding used:<br>○ Frequency of cage changes per week<br>*during gestation:*<br>*during neonatal period:*<br>*following weaning:*<br><br>d. Breeding - bred on site or timed pregnant, how many different sires (are the same fathers breeding with both experimental and control dams)<br>Breeding location: |

- Gestational age at shipping:
- Biological age of dams (if not listed in Section 8c):
- Number of Dams bred:
- How many times have dams been mated previously:
- How many times did the dams mate and not become pregnant:
- Are the dams primiparous or multiparous?
- What was the frequency of maternal handling during the gestational/neonatal period (e.g. cage cleanings, weighing, blood collection manipulations):
- Biological age of sires:
- Number of sires bred:
- How many times have sires been mated previously:
- How many times did the sires mate successfully (e.g. mating resulted in pregnancy, full term birth):
- If bred previously, what was the interval between mating times:
- Are sires matched to experimental and control dams:
- Describe the mating design (1:1, 1:2 etc):

e. Social enrichment – number of cage companions
   - Number of cage companions prior to breeding:
   - Gestational age when dam separated for parturition:
   - Number of cage companions at weaning:

f. Physical enrichment – describe enrichment devices, and when enrichment is in the cage (removed when pups born? Or present throughout study), does the enrichment type change? How frequently?
   - Describe what type of enrichment devices (and how many) are included in cage/housing room:

| | | |
|---|---|---|
| | | ○ Does enrichment type/access change across study?<br>○ If so, when does enrichment type/access change (e.g. enrichment removed prior to parturition and replaced in late neonatal period):<br><br>Additional Comments: |
| **Sample size**<br>➢Litter versus offspring<br><br>a. Specify the total number of animals used in each experiment, and the number of animals in each experimental group.<br>b. Explain how the number of animals was arrived at. Provide details of any sample size calculation used.<br>c. Indicate the number of independent replications of each experiment, if relevant. | 10 | Provide details of:<br>a. Maternal N vs offspring N<br>○ What is the total number of dams/litters included in the study:<br>○ What is the total number of offspring per litter included the study:<br><br>b. Litter size and sex distribution<br>○ What size was each litter maintained at:<br>○ What age did culling take place at:<br>○ How many males and females were maintained in each litter:<br><br>c. Cross fostering<br>○ Did cross fostering occur:<br>○ If so, at what age did cross fostering occur:<br><br>Additional Comments: |

| | | |
|---|---|---|
| **Allocating animals to experimental groups**<br><br>a. Give full details of how animals were allocated to experimental groups, including randomization or matching if done.<br>b. Describe the order in which the animals in the different experimental groups were treated and assessed. | 11 | a. How many offspring per litter were used in each measure:<br><br>b. Randomization/Matching procedures<br>○ What procedures were used to assign animals to groups:<br><br>c. Sex as a biological variable (behavioral and physiological outcomes)<br>○ Were both males and females evaluated in each behavioral and physiological outcome:<br><br>Additional Comments: |
| **Experimental outcomes**<br>➢Behavioral testing<br>➢Physiological endpoints<br><br>Clearly define the primary and secondary experimental outcomes assessed (e.g. cell death, molecular markers, behavioral changes). | 12 | a. Maternal behavior and pup interactions<br>○ If maternal care was evaluated, were there differences following immunogen challenge (if so, please briefly describe):<br><br>b. Age(s) of offspring at behavioral testing/physiological evaluation endpoints:<br><br>c. Order of testing (e.g. behavioral test order)<br>○ Were animals evaluated in a counter-balanced order in terms of:<br>*presentation of tests to each animal:*<br>*order of experimental/control groups run through each test:*<br>○ What was the inter-test interval if a single animal underwent a battery of tests: |

| | | |
|---|---|---|
| | | Additional Comments: |
| **Statistical methods**<br><br>a. Provide details of the statistical methods used for each analysis.<br>b. Specify the unit of analysis for each dataset (e.g. single animal, group of animals, single neuron).<br>c. Describe any methods used to assess whether the data met the assumptions of the statistical approach. | 13 | a. Unit of analysis for each data set<br>○ Is the unit (n) of each analysis based on number of litters, or number of animals used per group: |
| **Other Disclosures** | | Please make note of any other extraneous variables that you would like to report (e.g. fire alarms, construction, temporary relocations, other variables that you think we should be considering in our studies etc.): |

The recommended use of this reporting form is to fill it out and include it as supplemental material for each of your laboratory's research publications. If there are difficulties utilizing/adapting this fillable form, please contact one of the corresponding authors to request a copy. The authors give permission for this table to be edited for use in reporting on other animal models (e.g. postnatal immune challenge models, early life stress models) as appropriate.

Kentner AC, Bilbo AD, Brown AS, Hsiao EY, McAllister AK, Meyer U, Pearce BD, Pletnikov MV, Yolken RH, Bauman MD. (2018). Maternal immune activation: reporting guidelines to improve the rigor, reproducibility, and transparency of the model. Neuropsychopharmacology, https://doi.org/10.1038/s41386-018-0185-7.
